# Motor cortex facilitates visual action override through the superior colliculus

**DOI:** 10.64898/2026.11.14.766645

**Authors:** Kuisong Song, Giovanni Usseglio, Pierre Feugas, Sofia Morou, Peng Cui, Dimitris Mariatos-Metaxas, Ismael Navarro, Teresa Femenia, Theodoros Damoulas, Ramon Reig, Pierre Le Merre, Iakovos Lazaridis, Andreas A. Kardamakis

## Abstract

Unexpected visual events can leave goal pursuit undisturbed or trigger visual action override— an interruption that redirects behaviour towards the stimulus. Using a visual distraction task in freely moving mice, population calcium recordings and causal perturbations identified the lateral motor-related superior colliculus (lSCm) as a key node controlling distractor engagement. Retinal inputs to the sensory-related SC signalled distractors even when mice ignored them. Pre-distractor behaviour and motor cortical activity predicted subsequent engagement, while increased corticocollicular and decreased nigrocollicular activity before distractor onset were consistent with a permissive state for override. Suppressing corticocollicular transmission weakened override without detectably altering task engagement. Single-unit recordings in head-fixed mice revealed response amplification in subsets of lSCm neurons during combined retinal and motor cortical stimulation, despite weak responses to either input alone. Motor cortical input thus facilitates sensory recruitment of collicular action circuits, shaping the balance between action stability and action override during ongoing goal pursuit.

## Introduction

Our visual environment contains more information than can guide behaviour at any moment. Ongoing goals shape the selection of relevant information, in part through frontal cortical influences on visual processing^1–3^ . Yet events outside the current task can signal opportunities or threats that warrant immediate investigation^4,5^ . Adaptive behaviour therefore requires a balance between *action stability*, the maintenance of ongoing goal-directed behaviour despite competing sensory events, and *action override*, the interruption and redirection of that behaviour towards an unexpected stimulus. Such redirection can permit brief evaluation before the original goal is resumed, but can also constitute distraction from the task at hand. Although the sensorimotor circuits underlying orienting are well established, which neural circuits regulate susceptibility to behavioural interruption during ongoing goal pursuit remains unclear.

Preparatory activity in frontal motor networks supports goal-directed actions, while distributed cortical outputs can influence several downstream action circuits in parallel. The lateral motor-related superior colliculus (lSCm) provides a substrate for integrating these descending influences with sensory signals^6–8^ , receiving input from superficial visual layers, motor cortex and basal ganglia^9^ . Collicular circuits integrate spatial information across sensory modalities and generate orienting movements^10–13^ . They also participate in interactions with frontal cortex during the preparation and selection of instructed actions^14,15^ . Their responsiveness to visual events depends on the state of the motor system: in primate oculomotor tasks, preparatory activity modulates collicular visual sensitivity and the ability of distractors to redirect orienting^16,17^. These observations raise the possibility that motor cortical input regulates the readiness of collicular circuits to respond to competing sensory events during ongoing action. We reasoned that this input could establish a permissive collicular state that allows unexpected visual signals to recruit action circuits and interrupt goal pursuit.

To investigate this possibility, we developed a task in which freely moving mice initiated trials through visual search and navigated towards a visually cued reward. Visual distractors presented during navigation introduced an alternative target within the same sensory modality. Animals could continue towards the reward or engage with the distractor, providing a window onto the neural conditions governing action stability and override. Pre-distractor kinematics and motor cortical activity partially predicted subsequent engagement, suggesting that susceptibility depended on a state present before visual distractor onset. We combined simultaneous population calcium recordings with optogenetic and chemogenetic perturbations to investigate how descending corticocollicular inputs link this state to collicular recruitment. Complementary electrophysiology and modelling examined how cortical input facilitates visual recruitment of lSCm circuits, identifying circuit mechanisms that regulate susceptibility to action override.

## Results

### Susceptibility to visual action override varies with pre-distractor behavioural state

To examine how visual events interrupt ongoing goal-directed behaviour, we trained mice to locate a visual target in an open arena to initiate a trial (search phase), then navigate towards a visually cued reward port (goal phase). On 10% of trials, an off-axis visual distractor appeared to either side of the animal during the goal-directed trajectory (distractor phase; Fig. 1a). Across 574 trials from 10 wild-type mice, identical distractors produced outcomes ranging from pronounced interruption and reorientation to minor trajectory perturbations or uninterrupted navigation (Fig. 1b). Low-dimensional embedding of video-derived kinematic features (Fig. 1c) revealed a continuum of responses spanning four behavioural classes: *Neglect*, *Minor*, *Moderate* and *Strong* (Fig. 1d, Supplementary Video 2). On *Neglect* trials, mice continued towards the goal without overtly orienting towards the distractor. *Minor* engagement typically consisted of a brief head turn towards the distractor, either on the move or with a momentary pause. *Moderate* and *Strong* trials showed the clearest examples of action override. In both classes, mice stopped and oriented towards the distractor. On *Moderate* trials, they then resumed goal pursuit, whereas on *Strong* trials they spent several seconds exploring the distractor before returning to the task.

**Figure 1.**
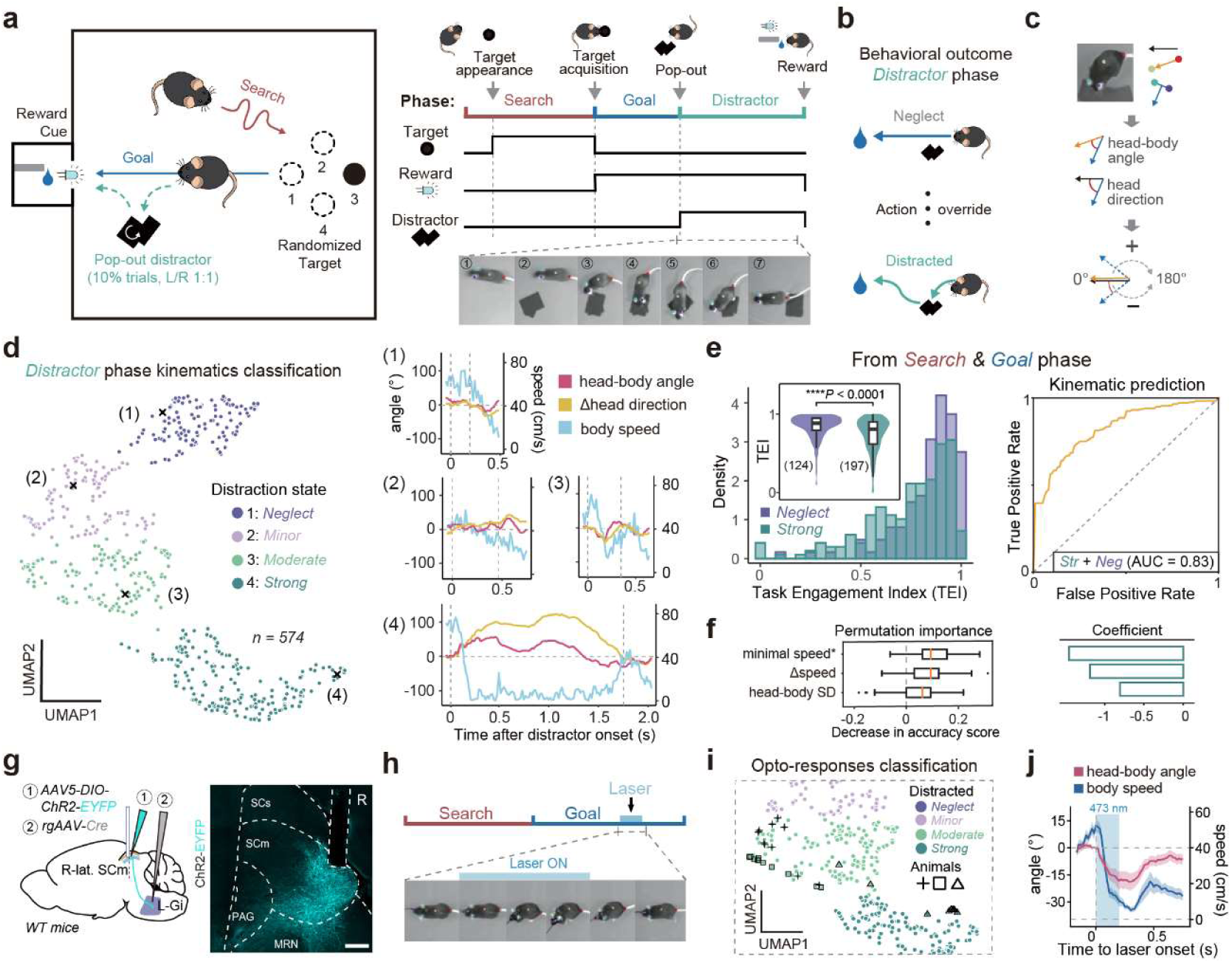
Pre-distractor behavioural state predicts visual action override, and activating medulla-projecting lSCm neurons mimics its kinematics. **a**, (*Left*) Behavioural task schematic. Trials start with a Search phase to acquire a randomized target (dark dot), followed by a Goal phase (navigation to the reward port). In 10% of trials, an interleaved Distractor phase presents visual distractors with equal lateral probability along the predicted trajectory. (*Right*) Time course of paradigm components in a visual distraction trial and a representative *strong* distraction response. **b**, Behavioural responses to distractors, ranging from neglect to *strong* engagement. **c**, Kinematic variable extraction. Body-part coordinates define head and body vectors to derive movement variables. Head direction is referenced to the horizontal axis, where angle signs denote lateral orientation (negative: left; positive: right) and absolute values represent deviation. **d,** (*Left*) UMAP embedding of visual distractor responses. Each point represents a response, color-coded by k-means cluster (*n* = 574 responses, *N* = 10 mice). (*Right*) Time series of movement variables for representative responses (1–4) from the clusters in panel d, color-coded as indicated. **e,** (*Left*) Task engagement index (TEI) distributions for *Strong* (dark green) and *Neglect* (dark purple) clusters. TEI integrates head trajectory and duration during Search and Goal phases, min-max scaled to 0–1 (low to high engagement). Inset: TEI comparison between clusters; sample sizes (n) in parentheses. Two-tailed Wilcoxon rank-sum test, \*\*\*\**P*<0.0001. (*Right*) ROC curves for logistic regression predicting distraction responses from pre-distraction kinematics, trained on *Strong* versus *Neglect* trials. AUC values are indicated in the legend. **f,** Permutation importance and coefficients of top 3 kinematic features for the model in panel e. **g,** (*Left*) Schematic of viral injection and fiber optic implantation in the right lateral superior colliculus motor-related layer (lSCm) and left gigantocellular nucleus (Gi) for optogenetic activation of lSCm-Gi projection neurons in WT mice. (*Right*) Representative coronal section of the SC; scale bar, 400 μm. White, ChR2-EYFP. **h,** Representative optogenetically evoked head re-orientation during the Goal phase of a no-distractor trial. **i,** UMAP embedding combining natural responses (from panel d) and optogenetically evoked responses. Optogenetic responses from individual animals are highlighted by distinct black symbols (*n* = 30 optogenetic stimulations, *N* = 3 mice). **j,** Average time series of head-body angle (red) and body speed (blue) of opto-evoked head re-orientation during the Goal phase (*n* = 30 opto-stimulations, *N* = 3 mice). Light blue shading indicates 473-nm laser stimulation. Data are mean ± s.e.m., vertical dashed line indicates laser onset (*t* = 0).

We next asked whether susceptibility to override could be predicted before distractor onset. A trial-wise task engagement index (TEI, see Methods), derived from path optimality and traversal time during the preceding search and goal phases, was higher before *Neglect* than *Strong* responses (*p*<0.0001; Fig. 1e, left). Pre-distractor kinematic features also discriminated *Strong* from *Neglect* trials, with an area under the receiver operating characteristic curve (AUC) of 0.83 (321 trials; Fig. 1e, right). The most informative features included the range and minimum of locomotor speed and variability in head–body angle during the goal phase (Fig. 1f). These predictive differences emerged while animals were pursuing the same goal, suggesting that susceptibility to interruption varies within goal-directed behaviour and need not reflect a transition into a distinct mode of exploration.

### Activating medulla-projecting lSCm orienting neurons mimics override kinematics

Given the role of the superior colliculus in visually guided orienting^18,19^ , we next asked whether recruitment of a defined collicular output pathway could reproduce the kinematics of override. Using a dual-vector viral strategy, we expressed channelrhodopsin-2 (ChR2)^20^ in glutamatergic lSCm neurons projecting to the contralateral gigantocellular nucleus^21,22^ (cGi; N=3; Fig. 1g). Optostimulation of these neurons, hereafter termed lSCm *orienting neurons*, during the goal phase interrupted locomotion and evoked rapid contralateral orienting, with transient stopping followed by head reorientation (Supplementary Video 3). When projected into the same behavioural space, stimulated trials fell predominantly within the *Moderate* and *Strong* response clusters (Fig. 1i). Activation of this pathway therefore reproduced key movement components of visual action override.

### Collicular recruitment scales with and regulates the strength of action override

Having established that activation of orienting neurons reproduces the kinematics of visual action override, we asked how endogenous lSCm population activity relates to its trial-to-trial variability. In vGAT-Cre mice, we simultaneously recorded population calcium dynamics using dual-colour fibre photometry^23^ (trials= 102, N=5; Fig. 2a). VGAT⁺ inhibitory neurons expressed Cre-dependent jRGECO1a, whereas a complementary Cre-OFF strategy drove GCaMP6s expression in a predominantly VGLUT2⁺ population (85.9 ± 0.02%, mean±SEM, in situ hybridisation; Extended Data Fig. 5d,e). We refer to these as inhibitory and excitatory populations, respectively. Recordings from the right lSCm allowed us to compare leftward movements with different behavioural functions: orienting towards a contralateral distractor interrupted goal pursuit (i.e. override), whereas a leftward return movement after engagement with an ipsilateral distractor redirected the animal towards the reward, restoring goal pursuit.

**Figure 2.**
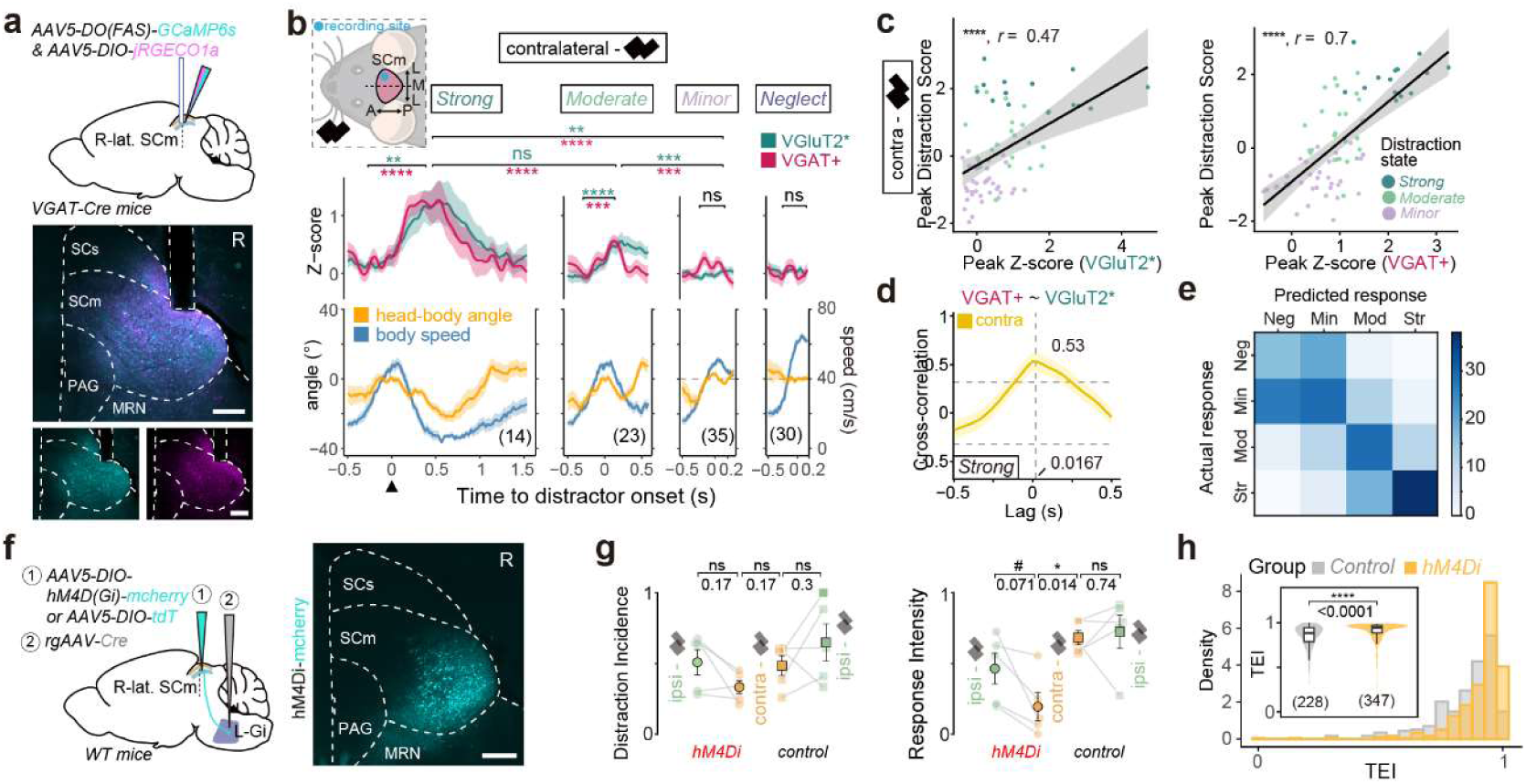
lSCm activity scales with the degree of distractor engagement, and chemoinhibition of medulla-projecting lSCm neurons weakens action override. **a,** (*Top*) Schematic of viral injection and fiber optic implantation in the right lSCm for simultaneous recording of putative GABAergic (Cre-ON) and glutamatergic (Cre-OFF) neurons in vGAT-Cre mice. (*Bottom*) Representative coronal sections of the SC; scale bar, 400 μm. Cyan, GCaMP6s; magenta, jRGECO1a. **b,** (*Top left*) Schematic of contralateral distractor placement relative to the recording site. Average time series of baseline-subtracted z-scored calcium signals (*Top*) and kinematic variables (*Bottom*) for responses to contralateral distractors, truncated at the median response duration. Total response counts (*n*) in parentheses. (*N* = 5 mice). Pre- and post-distractor peak comparisons (Two-tailed paired *t*-test, FDR-corrected) and cross-level post-distractor peak comparisons (Welch’s ANOVA with Games-Howell test) are indicated above traces. **c,** Pearson’s correlation between peak distraction score (DS, derived via PCA; see Methods) and peak z-scored calcium signals for putative VGluT2+ (VGluT2*) (*top*) and VGAT+ (*bottom*) neurons following contralateral distractors. Solid lines denote linear regression fits, shaded regions represent 95% confidence intervals (CIs). **d,** Cross-correlation analysis of VGluT2* (*x*) and VGAT+ (*y*) calcium signals during *Strong* responses for contralateral distractors. Vertical dashed lines indicate the lag at maximum correlation, horizontal dashed lines represent the 95% CIs determined by permutation testing (*n* = 1000 shuffles). **e,** Confusion matrix of logistic regression models predicting distraction levels based on SC calcium dynamics. Data are mean ± s.e.m., vertical dashed lines indicate distractor onset (*t* = 0) unless otherwise specified. \**P* < 0.05, \*\**P* < 0.01, \*\*\**P* < 0.001,\*\*\*\**P* < 0.0001, ns, not significant. **f,** (*Left*) Schematic of viral injection in the right lSCm and left Gi for chemogenetic inhibition of lSCm-Gi projection neurons in WT mice. (*Right*) Representative coronal section of the SC; scale bar, 400 μm. White, hM4Di-mCherry. **g,** Comparisons of distraction metrics across groups and distractor sides (*N* = 5 mice per group). Data represent mean ± s.e.m. (black outlines). Distractor sides are color-coded and groups are shape-coded. Two-tailed paired *t*-test (within-group) and two-tailed unpaired *t*-test (between-groups), FDR-corrected. #*P* (uncorrected) = 0.048 in g. **h,** TEI distributions for chemo-inhibition (orange) and control (gray, tdTomato + CNO) groups. Inset: comparison of TEI between groups, sample sizes (*n*) in parentheses. Two-tailed Wilcoxon rank-sum test.

We first analysed responses to contralateral distractors, retaining contraversive return movements from engaged ipsilateral trials as a control for movement direction (Extended Data Fig. 4c). Both excitatory and inhibitory populations were recruited during orienting, with larger responses accompanying higher degrees of distractor engagement (Fig. 2b). To quantify this, we derived a distraction score (DS) from instantaneous body velocity and head–body angle (Extended Data Fig. 4g). Peak activity in both populations correlated positively with DS (Fig. 2c), and their cross-correlation peaked near zero lag (Fig. 2d). Behavioural response classes could also be decoded from population activity, with misclassifications occurring mainly between adjacent engagement categories rather than between *Strong* and *Neglect* (Fig. 2e). Together, these findings link the recruitment of both excitatory and inhibitory lSCm populations to the degree of distractor engagement.

To test whether recruitment of the orienting neurons contributes causally to response strength, we unilaterally chemoinhibited orienting neurons using inhibitory DREADDs^24^ (*N*=5; Fig. 2f). We distinguished distraction incidence (DI), the proportion of *Strong* and *Moderate* responses among all distractor trials, from response intensity (RI), the proportion of *Strong* responses among *Strong* and *Moderate* trials (*see* Methods). We found that chemoinhibition reduced RI (*p*=0.014) without a detectable change in DI (*p*=0.17; Fig. 2g), shifting trials with moderate or *strong* engagement towards weaker classes, often terminating as brief head movements before reward-directed navigation resumed. Chemoinhibition was also associated with an increased TEI (*p*<0.0001). Thus, inhibiting the pathway reduced the degree of action override and favoured action stability; however, there was no evidence of a corresponding reduction in distraction incidence.

Inhibitory population activity closely mirrored excitatory recruitment, prompting us to test its causal contribution. Optogenetic activation of lSCm GABAergic neurons immediately before distractor onset usually attenuated contralateral reorientation towards the distractor (*N*=6; Fig. 3a-c) with ipsilateral turning biases or delayed engagement after stimulus presentation (Fig. 3d,e; *see* also Supplementary Video 4). Optostimulation alone did not detectably alter orientation before distractor onset. Distractor engagement was rarely abolished; its progression and execution were instead disrupted. Conversely, chemoinhibition of hM4Di-expressing GABAergic neurons across collicular layers (*N*=5), with injections targeted to the lateral SC, increased DI (*p*=0.025) without a detectable change in baseline task engagement (Fig. 3f-h). These perturbations implicate collicular inhibition in constraining distractor engagement, while the output-pathway manipulation identifies a contribution to response intensity.

**Figure 3.**
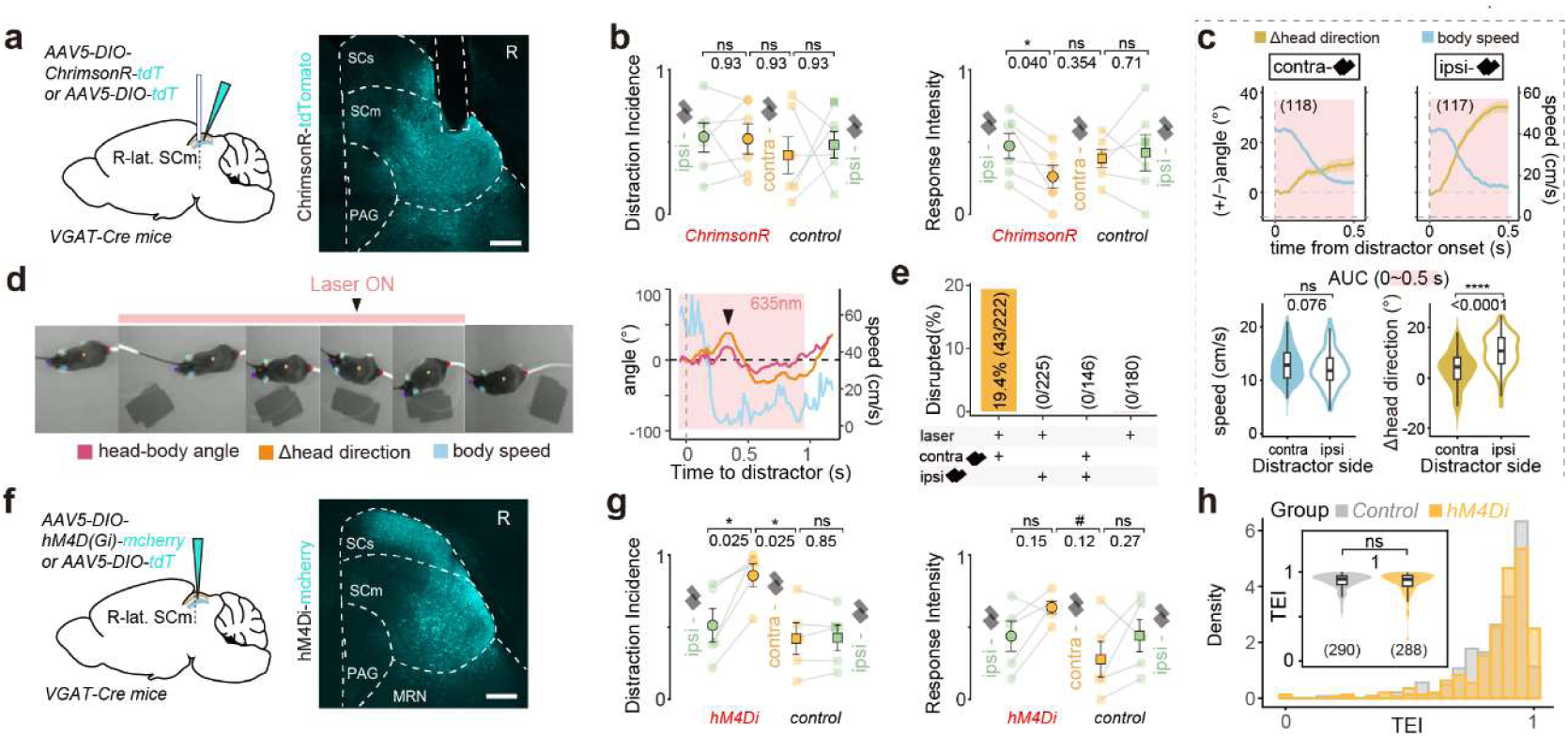
Activating collicular GABAergic neurons attenuates distractor-driven orienting, whereas inhibiting them increases the incidence of engagement. **a,** (*Left*) Schematic of viral injection and fiber optic implantation in the right lSCm for optogenetic activation of SC VGAT+ neurons in vGAT-Cre mice. (*Right*) Representative coronal section of the SC; scale bar, 400 μm. White, ChrimsonR-tdTomato. **b, g,** Comparisons of distraction metrics across groups and distractor sides (b, *N* = 6 mice per group; g, *N* = 5 mice per group). Data represent mean ± s.e.m. (black outlines). Distractor sides are color-coded and groups are shape-coded. Two-tailed paired *t*-test (within-group) and two-tailed unpaired *t*-test (between-groups), FDR-corrected. #*P*(uncorrected) = 0.039 in g. **c,** Kinematic analysis of post-distractor responses during laser stimulation. (*Top*) Average time series of Δhead direction (yellow) and body speed (blue) during the initial 0.5 s post-distractor onset. For contralateral distractors (*Left*), angular directions were negated for consistency with ipsilateral trials (*Right*). Red shading indicates the 0.5 s analysis window. (*Bottom*) AUC for body speed (*Left*) and Δhead direction (*Right*) during the 0.5 s window, comparing responses between contralateral (solid) and ipsilateral (hollow) distractors. Two-tailed Wilcoxon rank-sum test. **d,** Representative example of an optogenetically evoked disrupted distraction response. (*Left*) Chronologically ordered frames illustrate the behavioural progression. (*Right*) Time series of color-coded kinematic variables, the red shading indicates 635 nm laser stimulation. Arrows mark the initial counter-orienting (head deflection away from the distractor) preceding the subsequent re-orientation toward the distractor. **e,** Proportion of disrupted distraction responses across experimental conditions. Proportions and response counts (*n*) are displayed. **f,** (*Left*) Schematic of viral injection in the right lateral SC for chemogenetic inhibition of SC VGAT+ neurons in vGAT-Cre mice. (*Right*) Representative coronal section of SC; scale bar, 400 μm. White, hM4Di-mCherry. Data are mean ± s.e.m., vertical dashed lines indicate distractor onset (*t* = 0) unless otherwise specified. \**P* < 0.05, \*\**P* < 0.01, \*\*\**P* < 0.001, \*\*\*\**P* < 0.0001, ns, not significant. **h,** TEI distributions for chemo-inhibition (orange) and control (gray, tdTomato + CNO) groups. Inset: comparison of TEI between groups, sample sizes (*n*) in parentheses. Two-tailed Wilcoxon rank-sum test.

### Retinal distractor signals persist independently of behavioural outcome

The variability in lSCm recruitment raised the question of whether corresponding differences were already present in incoming visual signals. We therefore monitored the two major sources of visual input to the visual-related superior colliculus (SCs): the retina and visual cortex^25^ (Fig. 4a). Intravitreal injection drove GCaMP8s expression in contralateral retinal ganglion cells (RGCs), while jRGECO1a labelled ipsilateral visual cortical neurons projecting to SCs, enabling simultaneous recordings from the two inputs during behaviour (*N*=6; Fig. 4b,c). Retinocollicular projections exhibited an early calcium transient after distractor onset (50–150 ms), relative to no-distractor control trials (Fig. 4c). This response was present across behavioural classes, including Neglect (Fig. 4d), showing that the distractor was represented in retinal input even when it did not interrupt ongoing behaviour. Visual cortical projections to SCs showed prominent fluctuations throughout the task but no consistent distractor-evoked population response (Fig 4c; Extended Data Fig. 5g).

**Figure 4.**
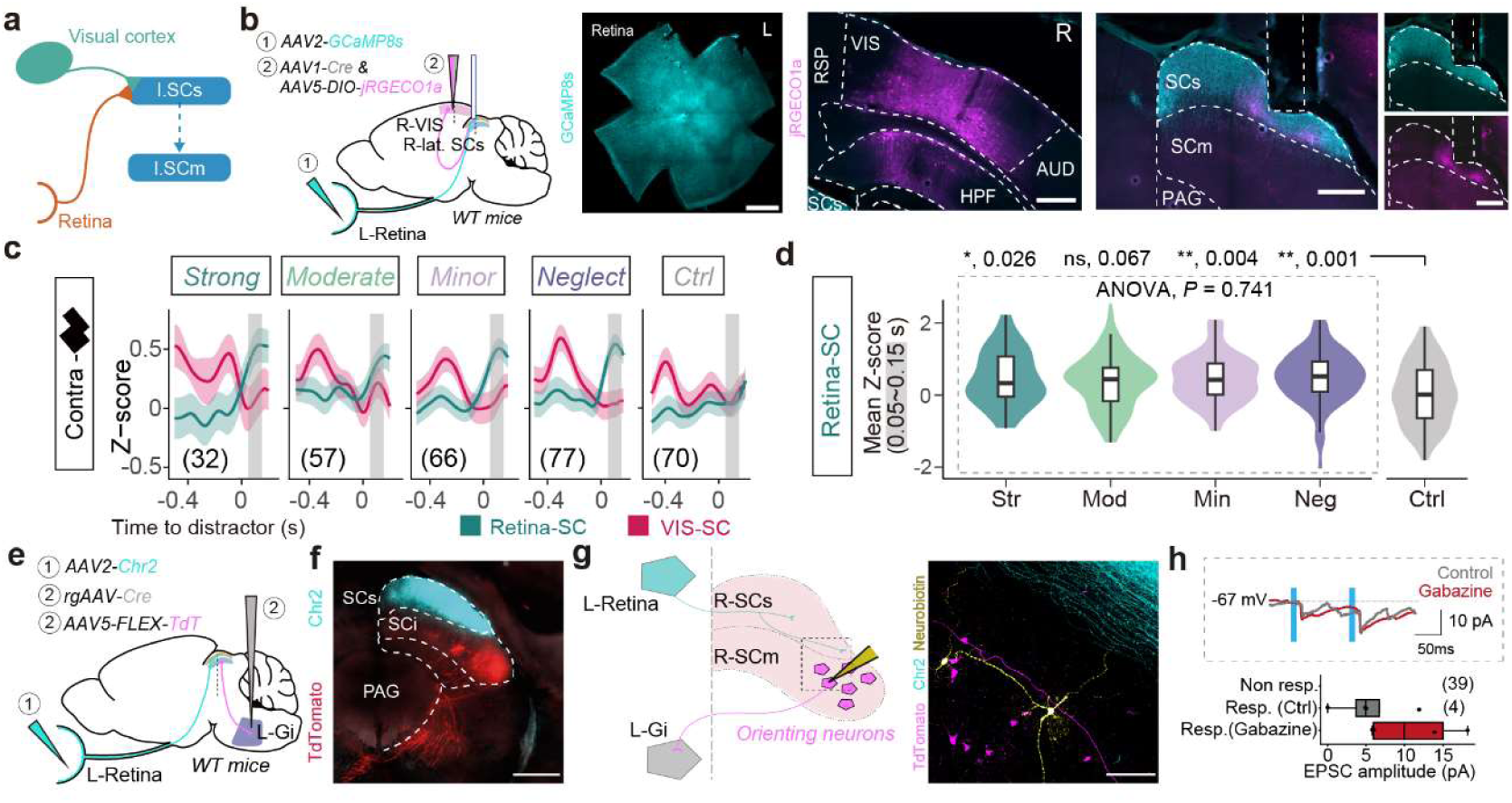
Retinal inputs signal distractors across behavioural outcomes, but retinal-terminal stimulation evokes little or no response in sampled lSCm orienting neurons. **a,** Potential circuit schematic underlying variable distraction responses. **b,** (*Top left*) Schematic of viral injection in the left retina and right visual areas (VIS), and fiber optic implantation in the right lateral superior colliculus sensory-related layer (lSCs) for simultaneous recording of Retina-SC and VIS-SC axonal terminals in WT mice. (*Bottom Left*) Representative retina; scale bar, 800 μm. (Right) Representative coronal sections of the VIS and SC (merged and individual channels); scale bar, 400 μm. Cyan, GCaMP8s; magenta, jRGECO1a. **c,** Average time series of z-scored calcium signals for Retina-SC (green) and VIS-SC (red) terminals across distraction levels and no distractor control. Gray shading (0.05–0.15 s) indicates the window for immediate response analysis. Total response counts (n) in parentheses. (N = 6 mice) **d**, Comparison of mean Retina-SC immediate dynamics during the shaded window across distraction levels compared to control, Welch’s ANOVA with Dunnett’s post hoc test. **e**, Schematic of viral injection in the left retina and left Gi for anatomical tracing of retinal projections and orienting neurons in the right SC. **f**, (Left) Representative coronal section of the SC. Cyan, Chr2. Red, TdTomato. (Right) Representative coronal section of the SC. Cyan, Chr2. Magenta, TdTomato. Yellow, Neurobiotin. **g**, (left) schematic of the patch-clamp recording set-up, the dotted inset represents the position of the image on the right. (right) Representative section of the SC. Cyan, Chr2. Red, TdTomato. Yellow, Neurobiotin. **h**. (Top) Representative trace of recorded neurons during light stimulation. (bottom) Box-plot of mean amplitudes of recorded lSCm orienting neurons after optogenetics stimulation of RGC terminals.

Preserved retinal input does not establish how effectively visual activity reaches lSCm orienting neurons. To examine this step, we expressed ChR2 in contralateral RGCs and performed whole-cell recordings from identified lSCm orienting neurons in midbrain slices (Fig.4e). Retinal axons densely innervated SCs, whereas the recorded orienting neurons lay deeper and their dendrites extended towards the the superficial neuropil (Fig. 4g). Most failed to respond (*n*=39) to optostimulation ChR2-expressing retinal terminals; whereas responsive cells yielded small amplitudes even in the presence of 20 μM gabazine (*n* = 4; mean EPSC amplitude: 5 ± 2 pA under control conditions and 11 ± 3 pA in the presence of gabazine; Fig. 4h) with inconsistent latencies spanning from 20 to 500 ms. These findings indicate weak effective retinal drive to the sampled lSCm output neurons under these recording conditions, despite dense retinal innervation of SCs. Given that the lSCm is a prominent target of motor cortical and nigral inputs, we next asked how these descending signals interact with visual input to regulate the recruitment of collicular orienting circuits.

### Cortical and nigral inputs regulate collicular susceptibility to visual action override

Motor cortical and nigral inputs provide routes for regulating collicular responsiveness. We first asked whether motor cortical activity before sensory arrival reflected susceptibility to override. Cortical surface electrocorticography (ECoG)^26^ at an interfacing point over the primary and secondary motor cortex (MOp/MOs) that give rise to dense collicular projections (*see* injection site) revealed differences between *Strong* (43 trials) and *Neglect* (67 trials) trials emerging approximately one second before stimulus onset (*N*=7; Fig. 5a), with power differing across all five frequency bands examined (α, β, γ, δ, θ; Fig. 5b). ECoG features discriminated Strong from Neglect trials with an AUC of 0.70 (Fig. 5c). Thus, motor cortical activity before distractor onset partly predicted subsequent action override, complementing the behavioural evidence that susceptibility to interruption is already reflected in the pre-distractor state.

**Figure 5.**
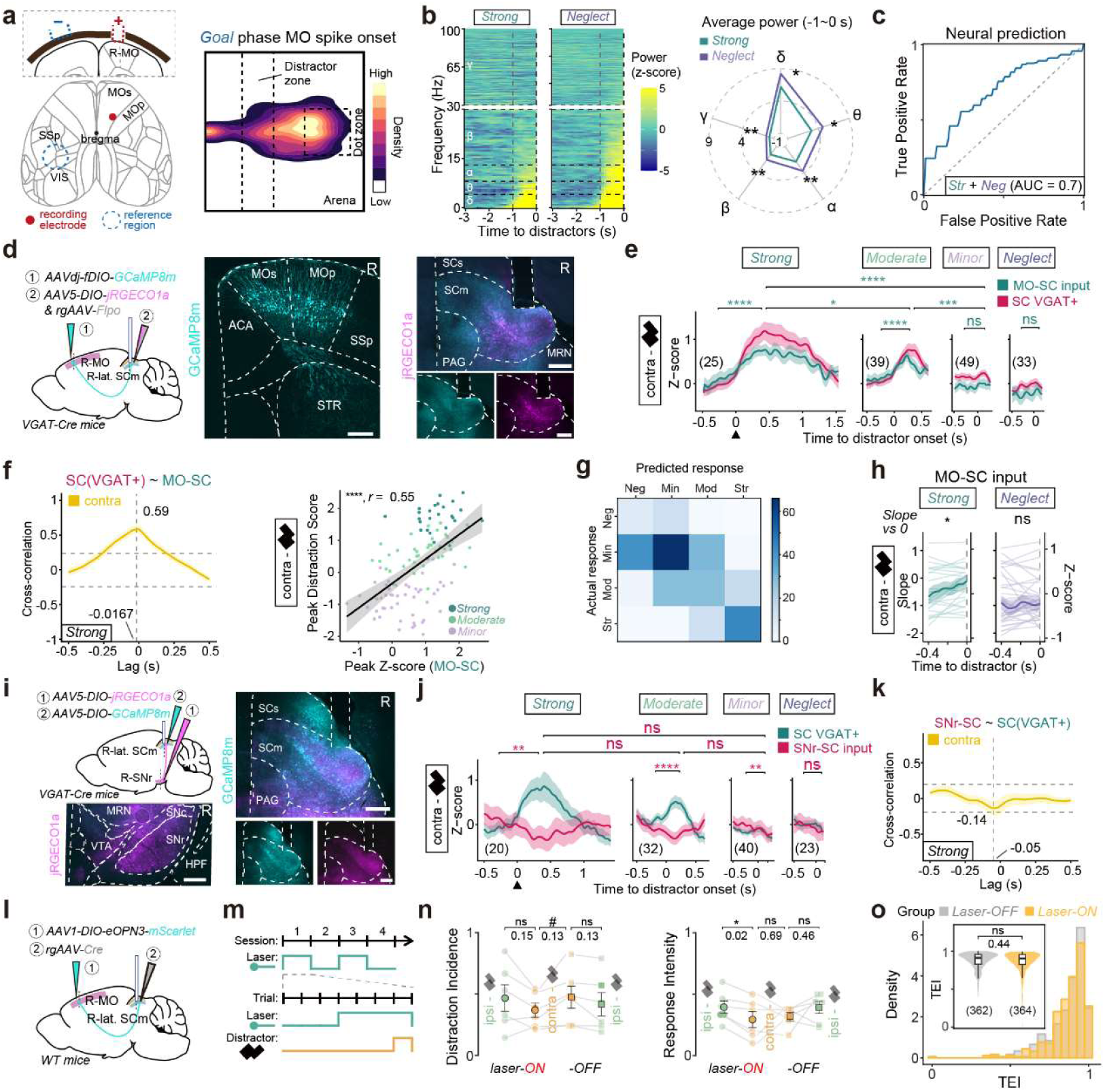
Motor cortical input to lSCm rises and SNr input falls before visual action override, and suppressing cortical input weakens override. **a,** (*Left*) Schematic of electrocorticogram (ECoG) electrode implantation in the primary and secondary motor cortex (MO; red, recording; blue, reference). (*Right*) Spatial density map of MO surface potential spike onsets during the goal phase. **b,** (*Left*) Average pre-distractor z-scored power spectral density (PSD) for *Strong* (*n* = 43 trials) and “Neglect” (*n* = 67 trials, *N* = 7 mice) responses. Dashed lines delineate pre-distractor window (-1 to 0 s) and frequency bands. (*Right*) Comparison of mean pre-distractor power between levels. Two-tailed *t*-test, FDR-corrected. **c,** ROC curves for logistic regression predicting distraction from pre-distractor MO spectral power (*Strong* versus *Neglect*). **d, i, l,** (*Left*) Schematic of viral injection and fiber optic implantation in right lSCm, MO or substantia nigra pars reticulata (SNr) for simultaneous recording of SC VGAT+ neurons and MO-SC (in d) or SNr-SC (in i) inputs in vGAT-Cre mice or opto-inhibition of MO-SC inputs (in l) in WT mice. (*Right*) Representative coronal sections of SC and MO (in d) or SNr (in i); scale bar, 400 μm. Cyan, GCaMP8m; magenta, jRGECO1a. **e, j,** Average time series of baseline-subtracted z-scored calcium signals for responses to contralateral distractors, truncated at the median response duration. Total response counts (*n*) in parentheses. (e, *N* = 6 mice; j, *N* = 5 mice). Statistical comparisons as in Fig. 2f. **f, k,** Cross-correlation of SC VGAT+ and MO-SC (*f left*) or SNr-SC (k) calcium signals during *Strong* responses for contralateral distractors. Dashed lines, lag at maximum correlation and 95% CIs. (*f Right*) Pearson’s correlation between peak DS and peak z-scored MO-SC signal following contralateral distractors. Solid lines, linear regression fits. Shading, 95% CIs. **g,** Confusion matrix of logistic regression predicting distraction levels from MO-SC input and SC VGAT+ dynamics. **h,** Pre-distractor MO-SC dynamics from e. Mean signal (line with shaded error) and per-trial linear fits (straight lines). Slopes were compared against zero, two-tailed one-sample *t*-test, FDR-corrected (\**P(Strong)* = 0.039). **m,** Schematic of laser delivery patterns across sessions and trials. **n,** Comparisons of distraction metrics across groups (Laser-ON/OFF) and distractor sides (*N* = 6 mice per group). Two-tailed paired *t*-test, FDR-corrected. #*P*(uncorrected) = 0.048. **o,** TEI distributions and comparison between groups, sample sizes (*n*) are in parentheses (*N* = 6 mice). Two-tailed Wilcoxon rank-sum test. Data are mean ± s.e.m., vertical dashed lines indicate distractor onset (*t* = 0) unless otherwise specified. \**P* < 0.05, \*\**P* < 0.01, \*\*\**P* < 0.001, \*\*\*\**P* < 0.0001, ns, not significant.

To examine the contribution of the descending pathway arising from this location, we simultaneously monitored calcium activity in lSCm-projecting MOp/MOs neurons projections and local VGAT⁺ lSCm neurons (N=6; Fig. 5d). The latter provided a measure of inhibitory population recruitment, which had covaried with excitatory recruitment in the separate recordings described above (Fig. 2). Corticocollicular and local inhibitory activity increased together during distraction and were strongly correlated (Fig. 5e,f). Corticocollicular activity scaled with response intensity and increased before distractor onset on *Moderate* and *Strong* trials; it was also used to decode behavioural response classes. In separate recordings of lSCm-projecting VGAT⁺ substantia nigra pars reticulata^27^ (SNr) neurons and local VGAT⁺ lSCm activity (*N*=5; Fig. 5i,j), nigrocollicular activity decreased before distractor onset on *Moderate* and *Strong* engagement trials, consistent with reduced nigral inhibition of the lSCm (Fig. 5k). Together with the pre-distractor increase in corticocollicular activity, this pattern is consistent with a permissive ‘two-key’ configuration in which cortical excitation and nigral disinhibition could jointly increase lSCm excitability, thereby favouring action override.

To determine whether the anticipatory corticocollicular activity observed during visual action override simply reflected preparation for contraversive movement (Fig. 5e), we examined task-directed movements in the same direction. Specifically, following engagement with ipsilateral distractors (Extended Data Fig. 7h), animals made contraversive return movements to resume their original goal. During these movements, corticocollicular activity accompanied lSCm recruitment and orienting, but lacked the anticipatory increase observed during distractor-driven override. Thus, although corticocollicular activity was recruited during both behaviours, its temporal relationship to movement differed depending on whether the movement interrupted or restored ongoing goal pursuit.

Building on previous work^28^, we next tested whether activating motor cortical input to lSCm was sufficient to evoke orienting (N = 5). ChR2-mediated stimulation of MOp/MOs terminals in lSCm across a range of intensities did not evoke orienting in an open arena without a visual distractor, despite histologically confirmed ChR2 expression (Extended Data Fig. 7m,n). Thus, corticocollicular activation alone was insufficient to elicit orienting under the conditions tested. Conversely, to suppress corticocollicular transmission during the visual distraction task, we expressed eOPN3^29^ in lSCm-projecting MOp/MOs neurons and delivered brief, intermittent green-light pulses to their terminals (0.1 Hz, 500 ms; for details see Fig. 5m) in lSCm, inducing prolonged presynaptic suppression. This shifted contralateral distractor responses towards weaker classes and reduced RI (P=0.02), without detectably affecting ongoing task engagement (N=6; Fig. 5l-o). Together, these findings support a permissive role – rather than an instructive one – for corticocollicular input in visual action override, facilitating distractor-driven orienting without being required for the maintenance of ongoing goal-directed behaviour.

### Coincident retinal and cortical activation reveals nonlinear collicular amplification

Given the causal contribution of corticocollicular input in visual action override and the weak effective retinal recruitment of lSCm output neurons, we reasoned that descending cortical input might facilitate the propagation and amplification of visual signals within lSCm. We therefore asked whether coincident activation of retinal and corticocollicular pathways could generate pronounced neuronal responses resembling the strong population recruitment observed during action override (Fig. 6a). To isolate these input interactions from overt orienting movements, we recorded single-unit activity^30,31^ in the lSCm of head-fixed mice (N=4, recording sessions=8, recorded units = 627) in head-fixed mice while optogenetically stimulating retinal and corticocollicular pathways separately or together (Fig. 6b–d). ChrimsonR was expressed in retinal ganglion cells and ChR2 in lSCm-projecting MOp/MOs neurons. Each stimulation alone evoked weak-to-moderate lSCm responses (37.6% ZETA test positive units during retinal stimulation, 31.4% ZETA test positive units during cortical stimulation). Combined stimulation (52% ZETA test positive units; see methods), however, revealed heterogeneous interactions, including early, delayed and sustained supralinear excitation, alongside sublinear responses and suppression (see methods; Fig.6e). Notably, subsets of lSCm neurons exhibited pronounced excitation absent during either input alone. Overall, population response during combined stimulation was significantly above the arithmetic sum of the response to individual stimulations (*p*<0.01; *see* methods; Fig. 6f). Thus, coincident activation of retinal and corticocollicular pathways generated nonlinear lSCm recruitment, providing a potential physiological basis for the *strong* collicular recruitment accompanying distractor engagement.

**Figure 6.**
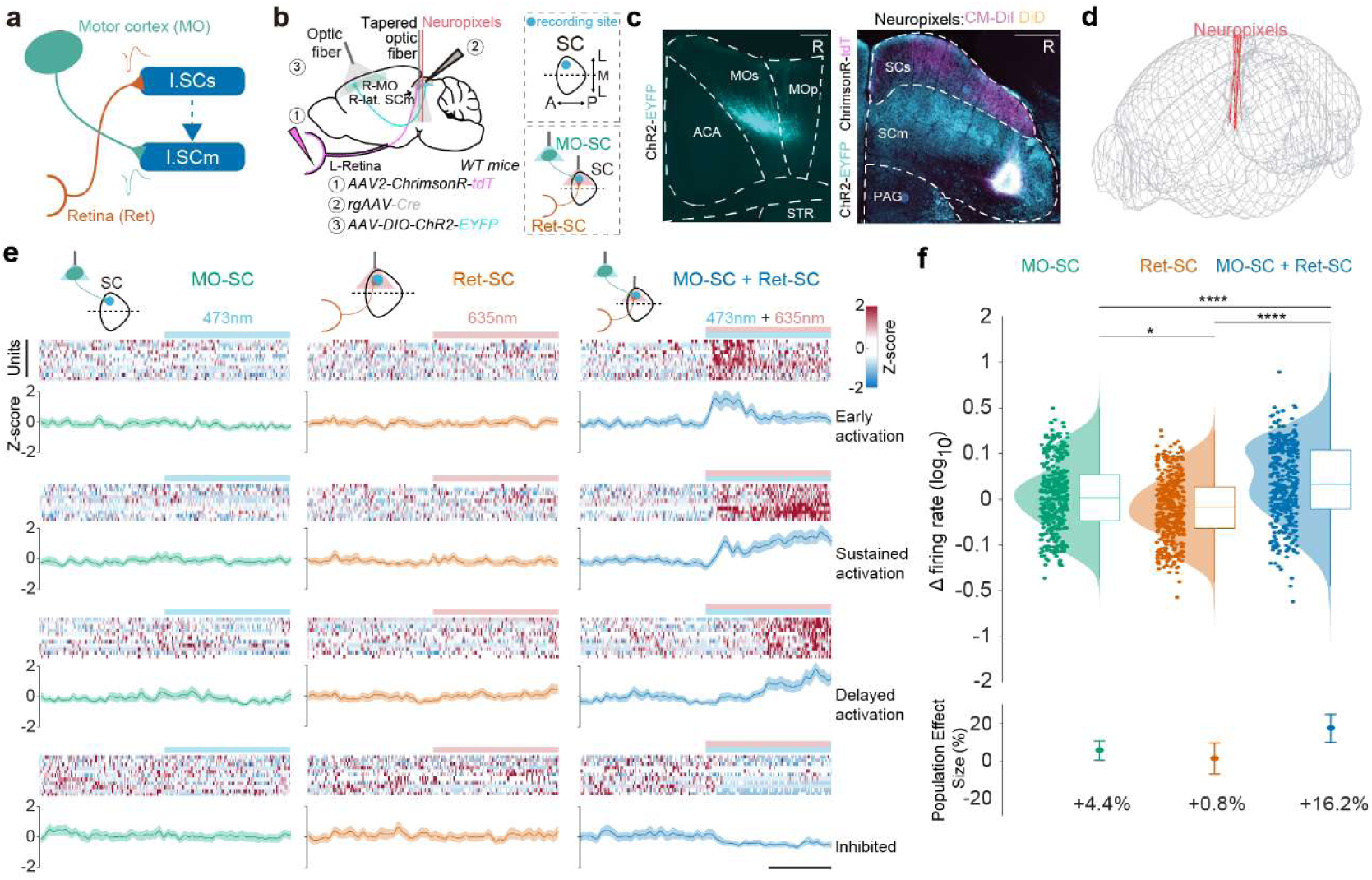
Coincident retinal and motor cortical stimulation evokes nonlinear responses in a subset of lSCm neurons. **a,** Potential circuit schematic underlying variable distraction responses. **b**, (*left*) Schematic representation of the experimental design of the viral injections in the superior colliculus (SC), in the contralateral retina and ipsilateral primary and secondary motor cortex (MO). An optic fiber is placed on the MO to stimulate cortical neurons and a tapered optic fiber is placed at the level of the SC to stimulate retinal axon terminals. A Neuropixels 1.0 probe is inserted at the level of the SC. (top *right*) Schematic of the unilateral recording site. (*bottom right*) Schematic of the optic fiber placement, Cortical (MO-SC) and Retinal (Ret-SC) optostimulation relative to the recording site. **c,** Representative histology of coronal slice of the MO (*left*) and SC (*right*); scale bar, 500 µm. **d,** Anatomical three-dimensional (3D) reconstruction of all probe tracks (N=4, recording sessions=8)**. e,** Representative Z-scored PSTH heatmap and Z-scored PSTH population average of single units recorded in the lSCm during three distinct optogenetic stimulation protocols. On the top of each column are represented schematics of the three lasers conditions. L*eft column,* stimulation with 473 nm to activate MO-SC inputs; *center column,* laser stimulation with 635 nm to activate Ret-SC inputs; right column, laser stimulation with 635 nm and 473 nm to activate MO-SC and Ret-SC inputs simultaneously. first row: early-activated units; second: sustained units; third: delayed-activated units; fourth: inhibited units. On the top of each column are represented schematics of the three lasers conditions. Vertical scalebar, 10 units. Horizontal scalebar, 500 ms. **f,** (*top*) Baseline-subtracted firing activity in Hertz (log^10^) of the three laser conditions shown in e (\**P* < 0.05, \*\**P* < 0.01, \*\*\**P* < 0.001, \*\*\*\**P* < 0.0001). (*bottom*) Population effect sizes for firing rate across the three laser conditions, expressed as a percentage change from the respective group baselines. Co-activation of retinal and cortical inputs was found to significantly increase SC firing rates by 16.2% versus baseline (P < 0.0001).

## Discussion

The same visual distractor could leave a mouse on course or interrupt its approach to the goal. Retinal inputs signalled the distractor in both cases, whereas recruitment of excitatory and inhibitory populations in the lateral motor-related superior colliculus scaled with the degree of distractor engagement. Perturbations identified lSCm as a key node in action override. Activating medulla-projecting lSCm neurons reproduced the stopping and reorientation characteristic of override, while other collicular manipulations altered the incidence or distractor engagement level. Suppressing motor corticocollicular input weakened distractor-driven orienting, whereas activating this input alone was insufficient to evoke orienting under the conditions tested. We therefore propose that susceptibility to action override depends on the collicular response to visual drive in the context of the preceding cortical state. Conceptually,

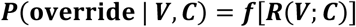

where **V** denotes visual drive, **C** the preceding corticocollicular activity, and **R(V;C)** the resulting lSCm response. In this framework, the same visual input can recruit orienting circuits to different degrees across trials, changing the likelihood that it interrupts goal pursuit.

Before the distractor appeared, movement kinematics and motor cortical oscillatory activity each partially predicted whether a trial would result in *Strong* engagement or *Neglect*. Greater oscillatory power was associated with action stability, whereas corticocollicular activity increased before distractor onset on both *Moderate* and *Strong* trials and scaled with subsequent engagement. These graded behavioural and neural conditions define what we mean by the pre-distractor state. To assess whether the pre-distractor cortical increase could be explained by movement direction alone, we compared two contraversive turns with different behavioural roles. One turn redirected the mouse towards a contralateral distractor. The other followed *Moderate* or *Strong* engagement with an ipsilateral distractor, as the mouse disengaged and turned back towards the goal. Both turns recruited corticocollicular and lSCm populations, but the rise in corticocollicular activity that preceded distractor-driven orienting had no comparable counterpart before the return to the goal.

Retinal inputs to the SCs reliably signalled distractors regardless of behavioural outcome, whereas visual cortical inputs showed no consistent distractor-evoked population response. These findings support a prominent subcortical role for retinotectal circuitry in registering unexpected visual events. The SCs can represent visual saliency even during visual cortical silencing and surround modulation by local circuits is conserved in the lamprey optic tectum and mouse superior colliculus^32–34^ . Within this conserved organisation, visual signals may reach orienting neurons by different routes. In lamprey, an ancient vertebrate lineage, deep tectal homologues of mammalian medulla-projecting orienting neurons receive direct retinal excitation through dendrites in superficial layers, yet inhibition dominates their visually evoked synaptic responses. In mice, by contrast, the dendrites of the sampled lSCm orienting neurons stopped short of the superficial retinal terminal field. This arrangement suggests that these neurons may receive weaker retinal-dependent drive than their lamprey homologues and rely more on di/oligosynaptic pathways through retinorecipient superficial interneurons^35^. The strength of visual drive from superficial layers to orienting circuits may therefore vary across vertebrate lineages and potentially across developmental stages, shaping how readily a salient event recruits an orienting response.

Motor cortical input provides one means of increasing the impact of weak visual drive. RGCs axonal terminal and corticocollicular optostimulation each evoked weak lSCm responses alone, yet coincident stimulation elicited nonlinear excitation and pronounced bursting in a subset of neurons. Ongoing modelling effort is exploring whether facilitation could begin near the interface between sensory- and motor-related collicular layers. Modest motor cortical excitation could enhance the recruitment of superficial glutamatergic neurons by retinal input, increasing visual drive to lSCm without requiring changes in interlaminar synaptic strength^34^. Narrow-field neurons,^36^, which project to deeper collicular layers, provide a candidate substrate for this relay. Nigrocollicular input may provide a complementary influence^27^. Before distractor onset on *Moderate* and *Strong* trials, corticocollicular activity increased and nigrocollicular activity decreased, with only the cortical increase scaling detectably with engagement. These separate recordings suggest a permissive ‘two-key’ configuration in which cortical excitation and reduced nigral inhibition could jointly increase lSCm excitability before sensory input arrives.

Lesion studies illustrate why the behavioural context of cortical control matters. In rats, motor cortical lesions prevented acquisition of a skilled movement sequence but spared its execution once learned^37^ . Human patients with unilateral frontal lesions could still orient towards a sudden visual target but had difficulty suppressing the reflexive saccade when instructed to look away. Such lesions affect multiple outputs, and their longer-term effects may include reorganisation of downstream circuits. During goal pursuit, selective suppression of motor cortical input to the lSCm weakened distractor-driven orienting without detectably altering task engagement. This dissociation suggests that maintaining an ongoing action and allowing a visual event to interrupt it may depend on partly distinct cortical outputs. By facilitating visual recruitment of collicular orienting circuits, motor cortical input may help explain how attentional capture finds expression in behaviour when an unexpected event briefly redirects action away from an ongoing goal.

## Acknowledgements

We thank Drs. Felix Leroy and Dimosthenis Pasadakis for their valuable input and support throughout this study. We also thank Dr. Cristina García Frigola for providing the viral vectors, and María Pérez Sanjuan for her technical assistance.

## Methods

### Animals and behavioural procedures

Experimental procedures were approved by the Committee on Animal Research at the Universidad Miguel Hernández and carried out in compliance with The Spanish National Research Council (CSIC), the Generalitat Valenciana (GVA) and European Regulations (2022/VSC/PEA/0236 to A.A.K.). Both male and female wild-type mice (C57BL/6N; Charles River) and VGAT-Cre mice (Slc32a1tm2(cre)Lowl/J, The Jackson Lab), aged 2∼6 months, were used throughout the experiments. During the behavioural training and experimental stages, mice were singly housed under a standard 12-h day/night cycle. Environmental conditions were maintained at a stable temperature (21 ± 1°C) and humidity (50–65%). The animals had *ad libitum* access to food, while water intake was restricted.

### Behavioural paradigm

Mice were trained to perform a goal-directed navigation task in an open-field arena. The behavioural sequence consisted of two phases: (1) Search phase: locating a visual target; (2) Goal phase: navigating to a reward port (hereafter “reward box”) and collecting reward, thereby completing one trial (Fig. 1a).

Each trial was initiated following reward collection from the previous trial. A dark circular target (4 cm diameter), displayed on a white background, served as the visual target. The target appeared 3-5 s after trial initiation and was positioned randomly at one of four predefined locations (25% probability per location) on the right side of the arena (Fig. 1a, Extended Data Fig. 1a). Once the mouse triggered the target by “entering” it, the target disappeared and the reward box, located on the left wall of the arena, was illuminated with white light as a “ready” visual cue. When the mouse performed a nose-poke into the reward box, a reward (8 μL of 15% sucrose solution) was delivered and the illumination ceased.

In visual distraction experiments, visual distractors were introduced in 10% of trials (typically every 7th∼10th trial), incorporating a distinct Distractor phase (Fig. 1a). Distractors were presented when two conditions were met: (1) the reward box was ready and (2) the mouse entered the left half of the arena. Distractors comprised two adjacent rotating dark rectangles (5-7 cm × 3 cm, angular velocity = 900°/s, clockwise or counter-clockwise rotation) positioned 6.5 cm lateral to the mouse’s expected trajectory. This predicted path was generated by linearly extrapolating the mouse’s real-time horizontal coordinates leftward from its position immediately before entering the left arena. Distractors appeared on either the left or right side of the predicted path with equal probability (50%) and remained visible until reward collection. Notably, in visual distraction trials, the target was exclusively presented at the rightmost position to promote a consistent horizontal trajectory toward the reward box.

### Training procedures

The training protocol comprised two sequential stages: (1) pre-training stage in an automated cage; (2) training stage in the arena. Mice were individually housed under water restriction throughout the entire training and experimental period, with food available *ad libitum*. Daily water ration (1-1.5 mL/day) was adjusted to maintain body weight at 90-95% (>85%) of baseline.

During pre-training, mice learned a classical nose-poke operant task in automated training cages. The behavioural sequence required initiating trials by nose-poking a designated port (Trial Initiation Box) on one chamber wall, followed by reward collection from a reward box on the opposite side (Extended Data Fig. 1c). Mice underwent daily 1-to 2-hours training sessions for six consecutive days, with multiple mice trained simultaneously in separate chambers (Extended Data Fig. 1e).

The training stage typically commenced after a 24- to 48-hour interval following pre-training. Mice were trained individually in the arena for 40-minute daily sessions, with duration adjustable based on performance. Most mice achieved proficiency by Day 4, defined as completing ≥80 trials within 30 minutes. Training sessions were scheduled at approximately the same circadian time as pre-training sessions.

### Behavioural rigs

The nosepoke box, a core component of our behavioural rigs, was 3D-printed (Ultimaker 2+) as a five-sided cube (L×W×H: 2.6×3×3 cm, thickness: 0.4 cm, open side: 3×3 cm), the bottom is removable for cleaning. One infrared (IR) LED (TSHF6410, Vishay Semiconductors) and one IR phototransistor (LTR-526AD, Lite-On) were mounted on two opposite sides of the box, near the opening, to detect nose-poking events. A white LED (LTW-420D7, Lite-On) was installed at the top to illuminate the nosepoke box as a “ready” visual cue. For the reward delivery in a reward box, an 18-G blunt needle was inserted through a rear hole, extending to the center to dispense sucrose solution (Extended Data Fig. 1d).

To build the automated training cage, one standard open cage (L×W×H: 32×18×13 cm) was divided into two chambers (L×W×H: 23×9×13 cm) with 3 Plexiglas boards. Each chamber contained a nosepoke box on opposing short walls, designated as the trial initiation box and reward box. Both boxes interfaced with an Arduino UNO Rev4 Minima microcontroller (ABX00080, Arduino) for automated control. Reward was precisely delivered using a mini peristaltic pump (3910, Adafruit), powered via a motor shield (1438, Adafruit). A thin silicone tube (ID: 0.5 mm, OD: 2.5 mm) was utilized to enable microliter-scale delivery. Each cage accommodated two mice simultaneously.

The arena (L×W×H: 45×45×32 cm) was constructed with black (walls) and transparent (floor) Plexiglas (thickness: 3 mm). Behavior was recorded using a Blackfly camera (bfs-u3-04s2m, FLIR) at 60 Hz (resolution: 720 × 540). A 50-inch screen (Hisense 50A6BG) beneath the arena displayed visual patterns on a white background. The arena was illuminated by two IR illuminators (CM-IR130-850NM, CMVision). A reward box mounted on the left wall delivered sucrose solution via a mini peristaltic pump (Model 80204, Campden Instruments) connected to an 18-G blunt needle. Pump and reward box control was managed by an Arduino Mega 2560 Rev3 (A000067, Arduino).

The closed-loop training program was developed using Bonsai and Arduino IDE, with Bonsai-Arduino communication achieved using an Arduino UNO rev3 board. The modular architecture included: video recording, mouse detection, mouse tracking, event trigger, event and frame timestamp recording, Arduino communication and visual pattern rendering (Extended Data Fig. 1b). Video acquisition relied on SpinnakerCapture Bonsai node (with SpinnakerSDK v 1.29.0.5); real-time mouse detection and tracking and detection were achieved with BackgroundSubtraction Bonsai node; Visual patterns were drawn by Bonsai BonVision package^41^. Bonsai training program and Arduino code are available upon request.

### Surgery

#### Viral injections

Injections were performed in WT or VGAT-Cre mice aged 2-6 months under 1-3% isoflurane anesthesia, with postoperative analgesia administered via subcutaneous Buprex (buprenorphine, 0.1 mg/kg). During procedures, mice were maintained at 37°C via heating pad with continuous ocular hydration (Viscotears gel). Mice were afforded a minimum 4-day post-operative recovery before behavioural training, and at least 14 days were allowed to ensure sufficient viral expression prior to recording or manipulation.

Adeno-associated viruses (AAVs) with titer ranging from 1×10¹² to 1×10¹³ viral genomes per milliliter (vg/mL) were utilized. For brain injections, viral suspensions were infused at 100 nL/min using a digital stereotaxic system (RWD 68528 frame with R-480 injector) through glass micropipettes (GC-3.5, RWD). Pipette was pulled by a puller (P-97, Sutter). The pulled micropipette was first positioned 0.1-0.3 mm beyond target depth to create viral dispersion space, then retracted to target coordinates for controlled infusion; After injection, the pipette was retracted by 0.1-0.3 mm and remained in position for 8 min to prevent backflow. For eye injections, Viral suspensions were delivered at 500 nL/min. A puncture was made 1-2 mm posterior to the corneal limbus using a sterile 30-gauge (G) needle after pupil dilation was achieved with 0.5% Alcaine. After which 1,000 nL vitreous was aspirated manually using a Hamilton microliter syringe with a 33-G blunt needle. The pulled micropipette was then inserted through the same puncture into the eye, and the tip was angled toward the vitreous humor and back of the eye, taking care to avoid any damage to the retina or lens. The pipette was connected to the injector with a thin silicone tube. After injection, the pipette remained in the eye for at least 30 s to prevent reflux of the viral suspensions.

The following viruses and coordinates were used for different experiments:

1. Fig. 2a, 4b, 5e S5e, a mix of 100 nL AAV5-CBA-DO(FAS)-GCaMP6s (Addgene plasmid #110135)^38^ and 100 nL AAV5-CAG-Flex-NES_jRGECO1a (Addgene plasmid # 100852) or 300 nL AAV5-hSyn1-dlox-ChrimsonR_tdTomato(rev)-dlox (v289-5, VVF UZH and Addgene plasmid #92207)^39^ or 300 nL AAV5-CBA-DO(FAS)-GCaMP6s only was injected into right lateral superior colliculus motor-related layer (lSCm) (ML 1.4 mm, AP -3.6 mm from bregma, DV 2 mm from brain surface) of VGAT-Cre mice.
2. Fig. 4b, 1 μL AAV2-syn-jGCaMP8s (Addgene plasmid # 162374)^40^ was injected into the left eye, at least 2 days after intravitreal injection, a mix of 200 nL AAV1-hSyn-Cre (Addgene viral prep # 105553-AAV1) and 200 nL AAV5-CAG-Flex-NES_jRGECO1a was injected into the right primary visual cortex (VIS) (ML 2.3 mm, AP -3.2 mm from bregma, DV 0.5 mm from brain surface) of WT mice.
3. Fig. 1g, 2f, 300 nL retroAAV-hEF1α-iCre (v225-retro, VVF UZH and Addgene plasmid #55636)^41^ was injected into the left gigantocellular nucleus (Gi) (ML 0.5 mm, AP -6 mm, DV 5.3 mm from bregma), 300-500 nL AAV5-hSyn1-dlox-hM4D(Gi)_mCherry(rev)-dlox (v84-5, VVF UZH) or AAV5-CAG-Flex-tdTomato (Boyden, UNC Vector Core) or AAV2-Ef1a-DIO-hChR2(H134R)-EYFP (Deisseroth, UNC Vector Core) was injected into right lSCm (same as (1)) of WT mice.
4. Fig. 3h, virus (same as Fig. 2A excluding retroAAV-Cre) was injected into right superior colliculus sensory-related layer (SCs)(ML 0.8 mm, AP -3.6 mm from bregma, DV 0.9 mm from brain surface) at a volume of 90 nL and lSCm (same as (1)) at a volume of 300 nL of VGAT-Cre mice.
5. Fig. 4b, 5d, 5e a mix of 100 nL AAV5-CAG-Flex-NES_jRGECO1a and 100 nL retroAAV-EF1α-Flpo (Addgene plasmid # 55637) or 200 nL retroAAV-hEF1α-iCre was injected into right lSCm (same as (1)), 300 nL AAV_DJ-hSyn1-chl-dFRT-jGCaMP8m(rev)-dFRT (v740-DJ, VVF UZH) or 300 nL AAV1-hSyn1-DIO-eOPN3_mScarlet (Addgene viral prep # 125713-AAV1)^29^ was injected into right frontal motor cortex (MO) (ML 1.1–1.3 mm, AP 1.1–1.42 mm from bregma, DV 0.8–0.9 mm from brain surface) of VGAT-Cre or WT mice.
6. Fig. 5i, 200 nL AAV5-syn-FLEX-jGCaMP8m (Addgene plasmid # 162378) was injected into right lSCm (same as (1)), 300 nL AAV5-CAG-Flex-NES_jRGECO1a was injected into right SNr (ML 1.5 mm, AP -3.2 mm from bregma, DV 4.3 mm from brain surface) of VGAT-Cre mice.
7. Extended Data Fig. 10a, for the rabies virus tracing, 300 nL retroAAV-hEF1α-iCre was injected into the left Gi (same as (3)), 400 nL helper virus AAV5-EF1a-DIO-TVA-V5-t2A-RG was injected into right lSCm (same as (1)) of WT mice. Three weeks later, 500 nL of the engineered rabies virus EnvA-coated G-deleted rabies virus with EGFP (EnvA-ΔG-Rb-EGFP) was injected into the same location at a 15° angle. The helper and Rabies viruses are gifts from Dr. Konstantinos Meletis at Karolinska Institutet.
8. Fig. 6b 1 μL AAV2-Syn-ChrimsonR-tdTomato (UNC Vector Core) was injected into the left eye, at least 2 days after intravitreal injection, 300 nL retroAAV-hEF1α-iCre was injected into right lSCm (ML 1.4 mm, AP -3.6 mm from bregma, DV 2 mm from brain surface) and a AAV2-Ef1a-DIO-hChR2(H134R)-EYFP (UNC Vector Core) was injected into right frontal MO (ML 1.1–1.3 mm, AP 1.1–1.42 mm from bregma, DV 0.8–0.9 mm from brain surface) of WT mice.
9. Fig 4e, 5g, 5h: 300 nL of ssAAV-retro/2-hEF1α-iCre-WPRE-bGHp(A) (retro) were injected into the left Gi, 300 nL of rAAV5-CAG-Flex-tdTomato were injected into the right lSCm (same as (1)) and 1 μL of AAV2-CAG-ChR2-GFP was intravitreally injected into the left retina.

#### Implantations

Immediately or approximately 10 days post-intracranial injection, a 200 μm (for optogenetic stimulation) or 400 μm (for fiber photometry recording) diameter fiber optic cannula (R-FOC-BL200C-39NA or R-FOC-BL400C-50NA, RWD) was stereotaxically implanted to the brain. Custom electrocorticogram (ECoG) electrodes were fabricated by soldering silver wires (AGW2010, Wire silver .50 mm, WPI) or stainless-steel screws (M1.0xL2.0 mm) (62513, RWD) to pin connectors (5482, WPI). To ensure robust adhesion of the implant, the skull surface was first mechanically roughened using a dental drill. A thin layer of OptiBond (25881E & 25882 E, Kerr) was then applied as a bonding agent, followed by the application of dental composite (595953VI, Tetric Evoflow, IVOCLAR VIVADENT) to construct a stable headstage. Both the OptiBond and the dental cement were light-cured using an ultraviolet (UV) lamp (Superlite1300, M+W Dental). The headstage was coated with black tape or epoxy resin to reduce light leakage. Mice were allowed a minimum 4-day recovery period prior to behavioural training.

The following coordinates were used for different experiments:

1. Fig. 5a, a silver wire was positioned on the surface of right motor cortex (ML 1.1 mm, AP 1.1 mm, DV 0.7 mm from bregma) for recording. A screw was anchored into the skull over the left-medial region without penetrating the underlying dura as reference electrode.
2. Fig. 1g, 2a, 3a, 5d, 5i, 5l, 6b a fiber optic cannula was implanted above the right lSCm. (ML 1.4 mm, AP -3.6 mm from bregma, DV 1.65–1.7 mm from brain surface).
3. Fig 4b, a fiber optic cannula was implanted above the right lSCs (ML 1.2 mm, AP -3.8 mm from bregma, DV 0.95 mm from brain surface) either perpendicularly or at a 10° lateral angle.

For electrophysiological recordings with Neuropixels probe, a lightweight metal head-post was secured to the skull using a combination of light-curing dental adhesive OptiBond and dental composite. For extracellular measurements, a recording chamber was established by constructing a dental cement perimeter around the target area of the skull.

### Imaging and manipulations

#### Fiber photometry and ECoG recording

After demonstrating consistent task performance, mice were connected to the patch cord during training sessions for at least two consecutive days to acclimate to the patch cord connection before experimental procedures. For secure patch cord connection, mice were briefly anesthetized (<1 min) with isoflurane, with behavioural tasks initiated no sooner than 5 minutes post-anesthesia to ensure full recovery.

Fiber photometry recordings were performed using a multi-channel photometry system (R821, RWD) controlled by official software. For simultaneous dual-color imaging of GCaMP and jRGECO1a, three LEDs (410 nm for calcium-independent isosbestic baseline, 470 nm for GCaMP, and 560 nm for jRGECO1a recording) were operated in an interleaved fashion at 60 Hz each (cumulative sampling rate of 180 Hz). The resulting fluorescence signals were captured by CMOS cameras. The photometry system was connected to a pigtailed rotary joint (400 μm core, 0.37NA, FRJ_1×1_PT_400-0.37_2m_FCM_0.15m_FCM, Doric Lenses), and then to the animal via a patch cord (400 μm core, 0.39NA, R-FC-L-N3-400-L1, RWD). Light intensity at the fiber tip was maintained between 20–50 μW, power was measured with a power meter (PM100D, THORLABS). To ensure precise synchronization between photometry and behavior recording, the behavioural camera was TTL-triggered by the photometry system.

ECoG signals were acquired using Spike2 software (Cambridge Electronic Design) at a sampling rate of 20 kHz. The recording and reference electrodes from the mouse were connected via flexible cables to a headstage pre-amplifier (CV-7B, Axon Instruments). The signals were then passed through a HumBug noise eliminator (A-M systems) before being amplified by an Axoclamp 2B amplifier (Axon Instruments). Final analog signals were digitized using a CED 1401 digitizer. For experimental synchronization, the TTL pulses for triggering the behavioural camera were simultaneously sent to the digitizer, ensuring aligned timestamps across systems.

#### Optogenetic manipulation

Optical stimulation was provided by 473 nm (MBL-FN-473-200mW, CNI laser), 532 nm (LRS-0532-GFM-00100-05, Laserglow), or 635 nm lasers (MRL-III-635L-100mW), which were coupled to the mouse through fiber-optic patch cords and a rotary joint. Laser pulses were TTL-triggered and modulated using a custom-programmed Arduino board. The following stimulation protocols were implemented:

1. Fig. 1g-j, ChR2-expressing orienting neurons in the lSCm were stimulated using single-pulse 473 nm laser (10 mW power at the tip, 190 ms duration). Laser was triggered through the same mode by replacing distractors in the experimental program with shortened interval, occurring in every 3rd trial.
2. Fig. 3a, ChrimsonR-expressing GABAergic neurons in the lSCm were stimulated using 635 nm laser (8mW power at the tip) in trains of 20 pulses at 20 Hz (10 ms duration). In the distraction trials, the laser and the distractor were co-triggered. Due to inherent circuit delay, the distractor onset followed the laser with a slight latency of 2–3 frames (equivalent to 33– 50 ms at 60 FPS). For the laser-only sessions, the laser replaced the distractor and was delivered every 3rd trial.
3. Fig. 5l-m, eOPN3-expressing MO-SC terminals in the lSCm were stimulated using 532 nm laser (8mW power at the tip) in trains of pulses at 0.1 Hz (500 ms duration). To accommodate the slow activation kinetics of eOPN3, photostimulation was initiated midway through the interval between two distraction trials. For instance, the laser train would commence approximately 3–5 trials prior to the next distractor and persist until the distraction trial concluded. This pre-activation strategy ensured that the opsin-mediated modulation reached a stable level during the distraction trials.
4. Extended Data Fig. 7m, lSCm-projecting MOp/MOs neurons were stimulated using single-pulse 473 nm laser (10 mW power at the tip, 190 ms duration). Laser was triggered through the same mode by replacing distractors in the experimental program with shortened interval, occurring in every 3rd trial.
5. Fig. 6, lSCm-projecting MOp/MOs neurons were stimulated using single-pulse 473 nm laser (1s). Axon terminals in the SC of ChrimsonR-expressing retinal ganglion cells were stimulates using single-pulse 635 nm laser (1s). Simultaneous stimulation was performed single-pulse 473 nm ansd 63f laser (1s).

#### Chemogenetic manipulation

Clozapine-N-oxide (CNO, 4936, Tocris) was administered to pharmacologically activate designer receptors exclusively activated by designer drugs (DREADDs). Specifically, the inhibitory hM4D(Gi) DREADD was used in the experiments. Stock solutions were initially prepared at 2 mg/mL and freshly diluted to 0.2 mg/mL prior to daily administration with 0.9% normal saline. Animals received intraperitoneal (i.p.) injections of 1 mg/kg CNO under brief isoflurane anesthesia (<1 min) approximately 10 minutes before behavioural testing. To minimize injection-induced stress effects, mice underwent a habituation protocol involving saline i.p. injections during 2 training sessions prior to experimental procedures.

### Electrophysiology

#### Whole-cell patch-clamp *ex vivo*

Whole-cell patch clamp recordings were performed on cGi-projecting tdTomato-expressing neurons of the lSCm, in WT mice at room temperature using pCLAMP 10 (Molecular Devices, USA). Midbrain slices were continuously perfused with oxygenated recording solution containing (in mM): NaCl 124, KCl 2.5, NaH₂PO₄(H₂O) 1.25, NaHCO₃ 26, Glucose 20, MgCl₂(6·H₂O) 1, CaCl₂(2·H₂O) 2 (pH 7.3 with HCl; ∼305 mOsm); at a rate of 2 mL/min during recording. The slices were visualized under an upright CleverScope motorized microscope (Micro Control Instruments, UK) equipped with a Dodt gradient contrast system (DGC-2), an oil condenser (U-AAC, Olympus, Japan), and two Chameleon3 cameras (Teledyne FLIR, USA). Whole-cell recordings were obtained under visual control using a 16× (0.80 NA) water dipping objective (CFI75 LWD 16X W, Nikon, Japan) combined with 2× c-mount extender (EX2C, Computar, USA) and c-mount adapters (U-TV1X-2 and U-CMAD3, Olympus, Japan) with Axon MultiClamp 700B amplifier and Axon Digidata 1550B digitizer (Molecular Devices, USA). Signals were sampled at 20 kHz. Pipettes used for recording were pulled from borosilicate glass capillaries (article no. 1403542, Hilgenberg, Germany) by a P-97 Micropipette Puller (Sutter Instrument, USA). Patch pipettes (5–8 MΩ) were filled with a Cesium-based intracellular solution contained (in mM): EGTA 2, HEPES 10, ATP-Mg 4, GTP-Na 0.3, phosphocreatine 10, NaCl 5, TEA-Cl 10, CsMeSO₃ 110 and QX314 3 (pH 7.2 with Cesium hydroxide; ∼290 mOsm). Due to the long recording times (usually 1 h), the access resistance was evaluated every 10 min to ensure that initial values (only under ∼30 MΩ were considered) did not increase throughout the recording session.

#### Head-fixed Neuropixels recordings *in vivo*

In the days preceding the electrophysiological recordings, mice underwent a 3- to 4-day habituation protocol. During this period, they were handled and gradually acclimated to head fixation by incrementally increasing the restriction duration from 15 minutes to 1 hour. A few hours before experimentation, two 300–500 µm craniotomies were opened at the targeted coordinates. Mice received isoflurane anesthesia and a subcutaneous injection of buprenorphine. After sealing the openings with Kwik-Cast (WPI), the mice were allowed to recover in their home cages.

To facilitate post hoc identification of recording sites, probes were coated with CM-DiI and DiD (Thermo Fisher). The coating was applied by repeatedly painting the probe shanks with a droplet of CM-DiI or DiD via micropipette; once dry, the shanks exhibited a distinct coloration. The reference electrode was first coupled to a silver wire for grounding placed on the pial surface. Using a micromanipulator (uMp-3, Sensapex), the probe was lowered at (speed ∼20 μ s^−1^) to a depth of 3,800 µm. A 20–30 minute stabilization period was observed before recording to allow the tissue to settle.

For optogenetics stimulation, an optic fiber was placed on the MO to stimulate cortical neurons and a tapered optic fiber^42,43^ (https://www.optogenix.com/lambda-fibers/; 0.39 NA/200 µm; active length: 3 mm; implant length: 30 mm; cLCF: φ1.25mm ceramic ferrule) was placed at the level of the SC to stimulate retinal axon terminals. The laser delivery was controlled via a custom-made script on a Raspberry Pi B 4GB. We recorded extracellular activity via Neuropixels probes (Phase 3B Option 1, IMEC; 383 sites over 3,800 µm). Data were digitized at 30 kHz (gain of 500) and transferred to a data acquisition system consisting of National Instruments PXIe modules (PXIe-1071, PCIe-8381, and PXIe-8381). We used SpikeGLX software for recording Neuropixels data.

Following the recordings, the probes were cleaned for 30 minutes in a fresh Tergazyme solution (Sigma-Aldrich) and subsequently rinsed with distilled water overnight.

### Histology

#### Immunofluorescent and *in situ* hybridization staining

Mice were anesthetized by pentobarbital (200mg/kg) and transcardially perfused with ice-cold 1× phosphate-buffered saline (PBS) and 4% paraformaldehyde (PFA). Brains were post-fixed in 4% PFA at 4°C overnight, then washed with 1× PBS 3 times. For immunostaining, the brains were trimmed and embedded with 4% agar, then coronally sectioned at 60 μm thickness using a vibratome (VT1000S, Leica). For *in situ* hybridization, brains were dehydrated in 15% sucrose for 24 h, followed by 30% sucrose for 24–72 h until they sank to the bottom. The tissue was then embedded in OCT compound (NEG-50, Thermo Scientific), frozen on dry ice, and stored at −20°C for short-term use. Coronal sections were obtained using a Leica cryostat (CM3050S, Leica Bio-systems) in 14 μm thickness. For retinal histology, intact eyeballs were enucleated following the aforementioned perfusion and collected simultaneously with the brain. The eyeballs were fixed overnight and washed with PBS. To prepare retinal flat-mounts, the retinas were dissected from the eye cups and flattened by making four radial incisions. The tissues were then washed three times in PBS.

For immunofluorescent staining, brain sections were incubated in primary antibody chicken anti-GFP (ab13970, Abcam, 1:2000) or rabbit anti-RFP (600-401-379, rockland, 1:2000) at room temperature (RT) overnight, then washed with 1× TBST (0.3% TritonX-100) 2 times for 10 min, and incubated in secondary antibody goat anti-chicken with AF488 (ab150173, abcam, 1:1000) or donkey anti-rabbit with Cy3 (711-165-152, Jackson Immunoresearch, 1:1000) at RT for 2∼4 hours, finally washed with 1x TBST (0.3% TritonX-100) 2 times for 10 min then 1x PBS 2 times for 15 min. All antibodies were diluted with 1× TBST (0.3% TritonX-100, 5% Donkey serum, 0.01% sodium azide). Retina staining followed the same protocol as described above. Brain sections and retinas were mounted with an antifade mounting medium (BSBT AR1109, VWR) and imaged using a Leica THUNDER Imager.

The fluorescence *in situ* hybridization was performed with RNAscope Multiplex Fluorescent Reagent Kit v2 (ACD, Cat. No. 323100). The probes for vGluT2 (Mm-Slc17a6-C2, ACD, Cat. No. 319171-C2) and vGAT (Mm-Slc32a1, ACD, Cat. No. 319191) were used to identify the cell types. The manufacturer’s protocol was followed. Immediately following the *in situ* hybridization, the brain sections were incubated in chicken anti-GFP (1:2000) antibody at 4°C overnight, then washed with 1× PBS and incubated in goat anti-chicken antibody with AF488 (1:1000). After PBS washing, the sections were mounted and imaged with the THUNDER Imager.

#### Neuropixels probe track reconstruction

Anatomical images containing the probe’s fluorescent DiI signal were registered to the Allen Institute Common Coordinate Framework (CCFv3). Probe trajectories were then estimated using the ‘SHARP-Track’ pipeline (https://github.com/cortex-lab/allenCCF). Based on the resulting orientation and position of the probe track, electrode locations were transformed into CCFv3 space. Individual units were assigned specific coordinates corresponding to the electrode site.

#### Quantification and statistical analysis

The following processing was carried out using custom R, Python or MATLAB scripts.

#### Behavioural video analysis

Mouse behavioural kinematics were quantified using DeepLabCut^44^ (version 2.3.9) through markerless tracking of four body landmarks: center of bilateral pinnae (as “left/right ear”), center of body (as “mid-spine”) and tail base (as “tail”). Directional vectors were mathematically derived as: (1) head vector: right to left ear; (2) body vector: mid-spine to tail. Three kinematic parameters were further computed: (1) head-body angle, defined as the angle between head and body vectors. (2) head direction, which is the angle between head vector and leftward horizontal axis. (3) body speed, derived from mid-spine landmark movement over time. Angular sign denotes lateral orientation (negative: left; positive: right), with the absolute magnitude representing the degree of deviation from the horizontal axis. (Fig. 1C).

#### Behavioural response categorization

As described in the behavioural paradigm part, experimental trials were partitioned into three phases: (1) Search phase; (2) Goal phase and (3) Distractor phase. For further analysis, we included only trials meeting two distractor-phase criteria: (1) with minimal distance to distractor centers between 1 and 9 cm during passage, and (2) absence of nondirectional exploratory behavior. Note that a typical hardware-induced latency of 50 ms (3 frames) between the trigger and distractor onset was compensated for by offsetting all distractor timestamps. Movement quantification focused on a behaviourally relevant temporal window within the distractor phase, spanning from distractor appearance to 300 ms (18 frames) before reward collection, excluding terminal period when mice typically passed distractors. From this time window, we derived 19 movement features across three kinematic parameters: (1) head-body angle and (2) change of head direction compared to the initial state (Δhead direction): maximal/mean/median angles, angular standard deviation (SD) and duration ratios exceeding 20° or 45° threshold ((angle > 20°)%, (angle > 45°)%); (3) body speed: minimal/mean/median speeds, speed SD, the difference between initial and minimal values (Δspeed) and duration ratio below 20% of the initial speed ((speed < 20%initial)%); Additionally, (4) behavioural window duration was also included (Extended Data Fig. 2a). After standard scaling, these features underwent dimensionality reduction through Uniform Manifold Approximation and Projection (UMAP), followed by k-means clustering (k=4) of the 2D embeddings to identify response patterns (Fig 1d).

To categorize responses from new mice based on the categorization in all the control mice, a logistic regression model was trained using the 19 movement features and 4 clustering classes of control group in distractor phase. The area under the curve (AUC) of the Receiver Operating Characteristic (ROC) curve was calculated based on the average validation results from 10-fold cross-validation. New responses were classified by loading movement features into this trained model.

To assess the contribution of each feature in the categorization model, we calculated the set of coefficients of all the features for each clustering class. The permutation importance of movement features was also computed, which, in brief, involves randomly shuffling the values of each features and measuring the decrease in classification accuracy; to mitigate the effects of multicollinear features, a hierarchical clustering based on the Spearman rank-order correlations was conducted in advance to reveal the correlations among features, a threshold of 0.1 was applied, only one representative feature was retained in every closely related feature cluster for further computation.

#### Behavioural Response Metrics

For the comparison of response levels towards left and right visual distractors, two ratios were computed:

1. Distraction Incidence (DI), percentage of distracted responses in all the responses, which is equal to the number of *strong* (cluster 4 in Fig. 1d) and moderate (cluster 3 in Fig. 1d) responses divided by the number of all responses.

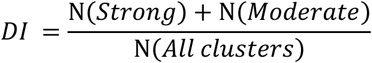

1. Response intensity (RI), percentage of *strong* responses in all the distracted responses, which is equal to the number of *strong* responses divided by the number of *strong* and *moderate* responses.

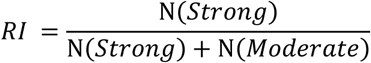

#### Task Engagement Index

To quantify the level of task engagement in mice, we developed a Task Engagement Index (TEI) based on the principle of behavioural efficiency (minimum energy expenditure). We defined high engagement as the animal’s ability to complete a trial using the shortest path and the fastest speed. The calculation focused exclusively on the Search and Goal phases to reflect the animal’s task-related state, intentionally excluding the Distractor phase to eliminate the influence of visual distractors. Movement trajectory was measured using mid-ears coordinates to capture both gross body movement and head turning. Duration and trajectory of Search and Goal phases were first Z-score standardized across trials to account for different units and scales, then these 4 standardized parameters were summed up, and applied a Min-Max normalization and inverted the values. The resulting TEI ranges from 0 to 1, where a value of 1 represents maximum task engagement (optimal efficiency) and 0 represents minimum engagement.

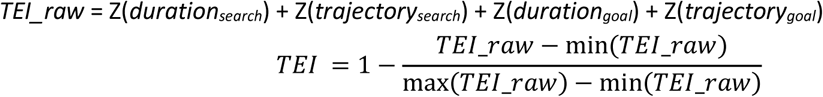

#### Response prediction and back-prediction

To investigate the possibility of predicting distraction response levels based on locomotion and reorientation features within the immediate prior period, 12 features were extracted from goal phase: the mean, median and SD of head-body angle and body speed, the angle between head direction and head-distractor direction (as head-distractor angle) right before response phase, the minimal speed, (speed < 20%initial)%, the maximal head-body angle and the duration of goal phase. Besides, we also included the duration of search phase, the immediate head direction and body speed of the mouse when it triggered the visual target. Based on these features, the prior behaviors were first clustered with UMAP to visualize the high-dimensional features and assess the separability of different trial types, every behavior was color-coded and shape-coded according to the response levels to distractors in the response phase of the same trial (Fig. 1F; Extended Data Fig. 3c). To predict responses from neural states, we extracted the mean power of standard frequency bands and the average local field potential during the 1-second window immediately preceding distractor onset. For the decoding of calcium indicator dynamics during the distractor phase, we derived 4 features for each channel, including the mean, median, peak amplitude, and SD.

A logistic regression model was then generated for the prediction using data from all trials or only trials with *strong* and *neglect* responses. The effects of multicollinear features were mitigated as mentioned before. To evaluate the performance, the AUC of the ROC curve was calculated based on the average validation results from 10-fold cross-validation. In small sample size conditions, repeated stratified 5-fold cross validation (10 repeats) was used instead (Fig 1L), and features were selected with univariate analysis (ANOVA F-test) by retaining only significant features to mitigate the risk of overfitting. For models integrating motion features with ECoG or calcium indicators, a feature pre-selection was employed to reduce dimensions. Specifically, we first conducted feature permutation importance analysis within each data modality independently. The top-ranked features from each group were then extracted and integrated into a multi-modal feature set. Comparative performance evaluations between models were subsequently performed using only these optimized feature subsets.

#### ECoG data analysis

The raw ECoG^26^ signals were initially sampled at 20000 Hz. To reduce computational complexity while maintaining relevant physiological information, signals were resampled to a sampling rate of 1000 Hz using a polyphase decimation filter. To eliminate power-line interference, a notch filter was applied at 50 Hz using a second-order IIR notch filter with a quality factor Q of 30. Zero-phase filtering was also implemented to prevent phase distortion. Power spectral density (PSD) was estimated using Welch’s method. Preprocessed ECoG signals were divided into overlapping segments using a 2-second Hamming window with 95% overlap. For each segment, the Fast Fourier Transform (FFT) was computed and averaged to obtain a robust power spectrum (0–100 Hz). The resulting absolute power values were then log-transformed into decibels (dB) and further standardized into Z-scores. The processed signals were decomposed into five standard physiological frequency bands: Delta (1–4 Hz), Theta (4–8 Hz), Alpha (8–13 Hz), Beta (13–30 Hz) and Gamma (30–80 Hz). Synchronization with behavioural videos was achieved by subtracting the recorded timestamp of trigger signal from raw timestamps.

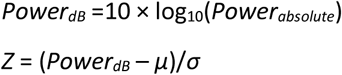

#### Fiber photometry data analysis

Python script was adapted from published work^23^ . Raw fluorescence signals were recorded from two experimental channels (560 nm for jRGECO1a or 470 nm for GCaMP) and one isosbestic control channel (410 nm). The isosbestic channel was used to account for motion artifacts and photobleaching. To reduce high-frequency noise while preserving signal integrity, both the experimental and control signals were subjected to low-pass filtering. We employed a 2nd-order Butterworth filter with a cutoff frequency of 4 Hz. Zero-phase filtering was implemented to prevent phase distortion and ensure temporal alignment of the signals. Photobleaching was corrected by fitting a double exponential function to the denoised signals. This fitted baseline was then subtracted from the denoised data to produce detrended signals. Movement-related artifacts were removed by leveraging the correlation between the isosbestic control channel and the experimental channel. A linear regression was performed to model the relationship between the detrended control channel and the detrended experimental channel. Then the estimated motion component was calculated by applying the linear regression model on the detrended control signal. The final motion-corrected signal was obtained by subtracting the estimated motion from the detrended signal. To allow for comparisons across different animals and trials, the corrected signals were normalized using Z-score, by subtracting the means and dividing by the standard deviations.

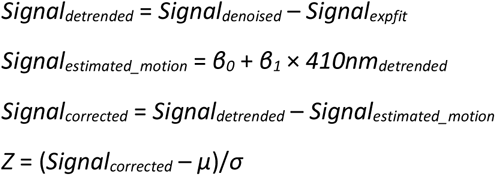

To isolate distractor-specific responses from background task activity, calcium signals from the preceding non-distraction trials were used as controls. We calculated the mean activity around the simulated “distractor” onset and applied time-proportional stretching to match the duration of each response. This estimated control signal was then subtracted from the distraction-trial data within the analysis window to account for task-related baseline fluctuations.

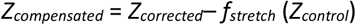

where *f_stretch_* denotes the duration-proportional temporal rescaling.

#### Cross-correlation analysis

To quantify the temporal relationship and correlation between the two calcium signals, we performed a Cross-Correlation Function (CCF) analysis. For the observation window (ranging from 400 ms prior to distractor onset to the median distractor phase duration) of each trial, Pearson cross-correlation coefficients were calculated between the two signals across a range of lags (up to ±30 points). A grand average CCF curve was generated by calculating the mean and standard error of the correlation coefficients across all responses at each lag. Statistical significance was determined using a global permutation test (N = 1000 shuffles). For each trial, the second signal was circularly shifted by a random interval (minimum 10% of signal length) to disrupt temporal alignment while preserving its original autocorrelation structure. A null distribution was constructed from the maximum absolute values of the resulting surrogate grand average CCF curves. The 95% global confidence threshold was defined as the 95th percentile of this distribution; any observed CCF peaks exceeding this threshold were considered significant at *P* < 0.05. CCF analysis was performed using the stats::ccf in R, following the convention *r*(*lag*) = cor(*x_t_*, *y_t-lag_*]), a positive lag indicates that signal *y* precedes signal *x*.

#### Distraction Score

A Distraction Score (DS) was developed using Principal Component Analysis (PCA) to integrate speed and head-body angle during Distractor phase into a unified metric. Variables were first adjusted so that higher values consistently represented increased distraction: speed was negated (–speed) to reflect slowing down during distraction; left head turning, which was reflected by negative head-body angles, was also negated (–head-body angle) to align with the right side turning. The data were then Z-score standardized to account for differing units. PCA was then performed on the scaled dataset of all trials. The first principal component (PC1) was extracted as the DS, as it captured the maximum shared variance between the two behavioural indicators. To ensure the index was intuitive (i.e., higher DS = higher distraction), the sign of PC1 was automatically flipped if it showed a negative correlation with the original distraction indicators (e.g. –speed).

#### High density electrophysiological data processing

The high-pass-filtered spike-band data underwent preprocessing via common-average referencing. First, each channel’s median was subtracted to eliminate baseline offset fluctuations. Subsequently, the median across all channels was subtracted from each individual channel to suppress shared noise and artifacts.

Data were spike-sorted with Kilosort 4.0, followed by manual curation using phy2. We excluded clusters characterized by non-physiological shapes, low amplitudes, or refractory period violations. Surviving units were then compared with adjacent clusters; merges were performed if units showed matching waveforms and synchronized refractory periods. A merge was finalized only if the resulting cluster demonstrated a clear refractory period and stable waveform consistency.

To generate the Z-scored PSTH heatmap and Z-scored PSTH population average, we binned the trial-average spike trains into 10 ms intervals and, in the case of population average, applied a Gaussian kernel for smoothing (Sigma = 16 ms). For statistical analysis, a quality control step was first performed, and units with firing rate < 0.5 Hz were discarded. To determine the responsiveness of single units, we employed the ZETA test, a parameter-free method for detecting statistically significant changes in spiking activity ^45,46^. Only units with significant ZETA test response in at least one laser protocol, were selected for analysis (391 ZETA test positive units selected in total). For each unit, trial-averaged firing rates were calculated over a 1-s laser presentation window and a 1-s pre-laser baseline period. To assess the effect of laser stimulation while accounting for the hierarchical structure of the data and inter-animal variability, we fitted a linear mixed-effects (LME) model. Firing rates were log-transformed to meet model assumptions of normality and homoscedasticity (Extended Data Fig. 10 c,d), with the difference between the log-transformed laser and baseline firing rates serving as the response variable. Laser condition was included as a fixed effect, while random intercepts were specified for animal and for individual units nested within animals to capture baseline variability and account for within-subject unit correlations. To return the fixed-effect estimates (beta) to the linear scale, beta values were exponentiated (10^beta) to derive the geometric mean fold change. Percentage changes in population firing rate were computed as (10^beta - 1) * 100. Error bars represent back-transformed 95% confidence intervals ((10^CI – 1) * 100), computed on the log scale by MATLAB’s “fitlme” using parameter standard errors and a Student’s t-distribution based on the model’s degrees of freedom. To evaluate differences between laser responses and test for supralinearity, we performed post-hoc linear contrasts using the Wald test on the fixed-effects coefficients of the linear mixed-effects model. P-values were corrected for multiple comparisons using Bonferroni method.

