## Extended Data for "Motor cortex facilitates visual action override through the superior colliculus"

### Motor cortex sets collicular state for visual action override

### Extended Data

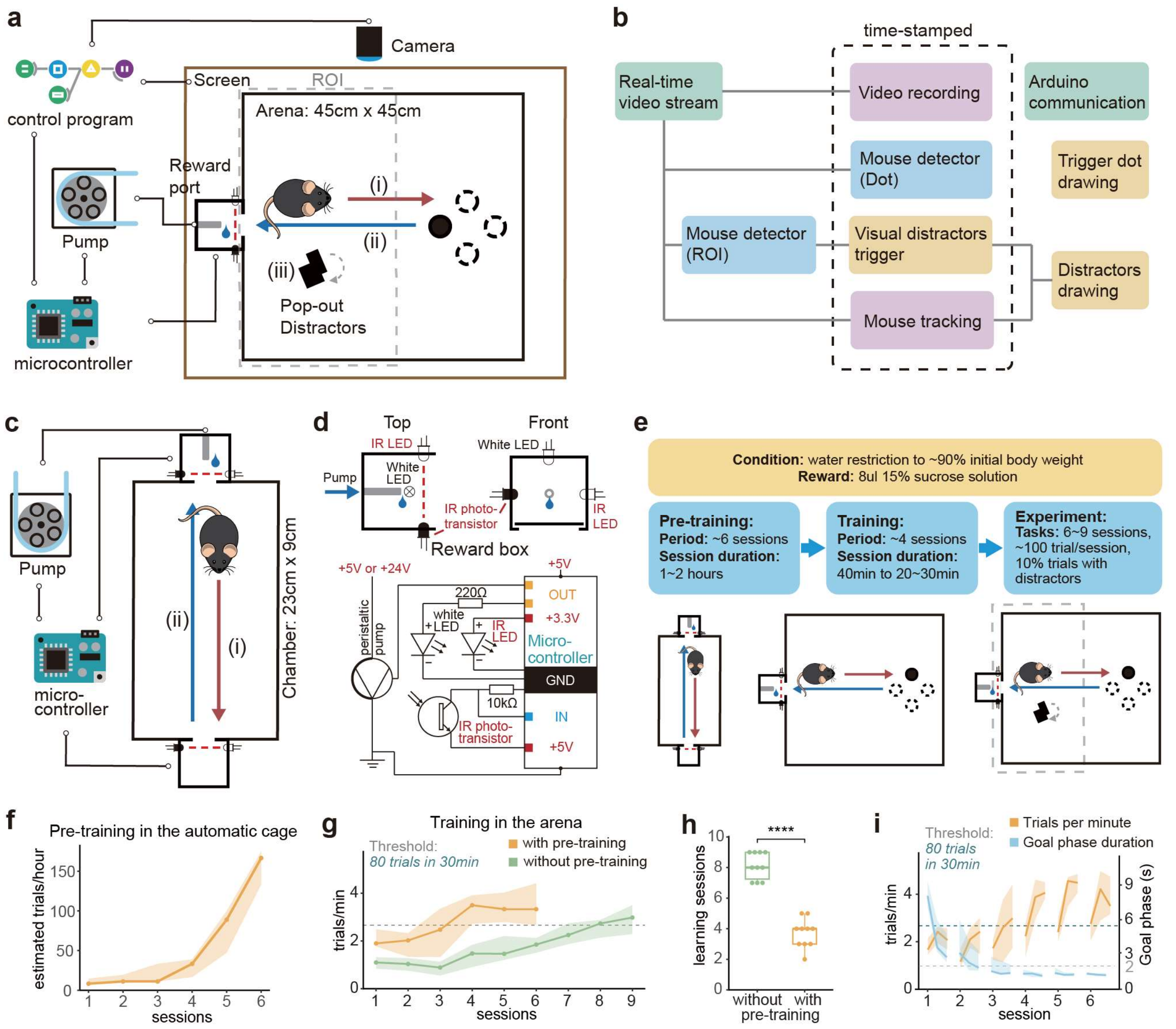


**Extended Data Fig. 1: Automated platforms and optimized strategies for efficient training of the visual distraction paradigm**

**a,** Schematic of the behavioral paradigm and experimental setup. A complete visual distraction trial consists of three sequential phases: (i) Search, (ii) Goal, and (iii) Distractor. The closed-loop system is fully automated via a custom control program and microcontrollers. The reward port detects nose pokes using infrared beam-break sensors. Visual stimuli (target and distractors) are displayed on a screen positioned beneath the arena. Dashed grey box labeled left half of the arena. **b,** Modular organization of the control program. Block diagram illustrates the functional components of the visual distraction task and their interconnections. **c,** Schematic of the Automated pre-training setup. Mice are first trained to trigger a nose-poke (i) and subsequently return to the port to collect a reward (ii) in a closed-loop manner. **d,** Hardware and electrical specifications. (*Top*) Top and front views of the reward port. (Bottom) Electrical circuit diagram of the automated cage. Constant voltage outputs (3.3 V or 5 V) and logic-controlled 5 V outputs are highlighted in red and orange, respectively. **e,** Experimental timeline. Mice were maintained under continuous water restriction. The procedure includes 6 sessions of pre-training, approximately 4 sessions of basic task training (without distractors), and the final experimental stage with visual distractors. One session per day. **f,** Learning progress during pre-training ($N$ = 10 mice). **g,** Learning curves of training stage with (orange) or without (green) pre-training.the horizontal dashed line at 2.67 trials per minute (80 trials per 30 minutes) marks the threshold for being well-trained. ($N$ = 10 mice per group). **h,** Comparison of number of sessions required to reach the training threshold with (orange) or without (green) pre-training. Two-tailed $t$-test ($N$ = 10 mice per group, ****$P < 0.0001$). **i,** Detailed behavioral dynamics during the training stage. Each session is divided into three equal segments. The orange line shows trial frequency (trials per minute), and the blue line shows Goal phase duration. ($N$ = 10 mice). Horizontal dashed lines indicate the 2.67 trials/min and 2 s duration thresholds. Solid line and shaded area represent median and IQR in panel f, g, h.

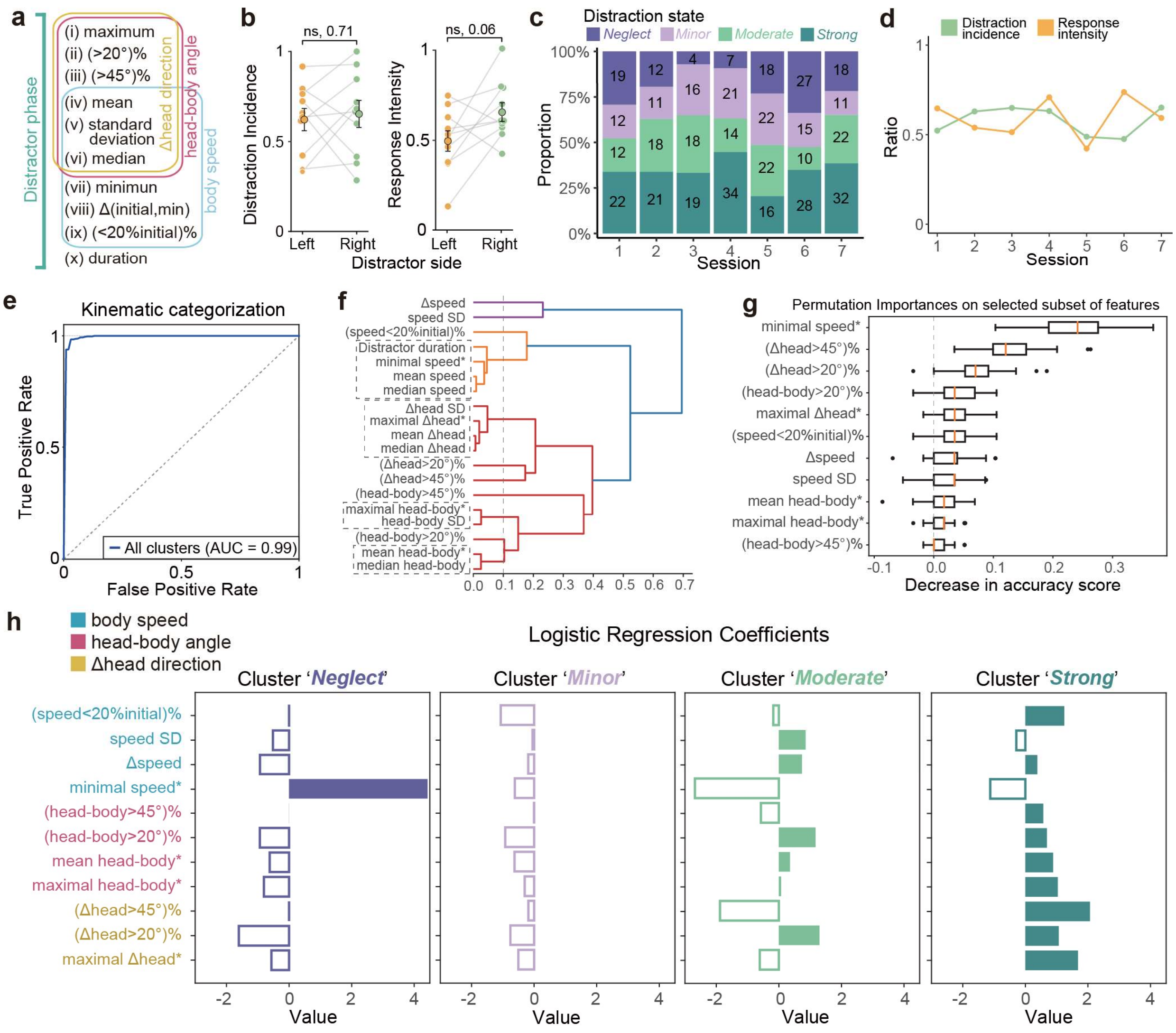


**Extended Data Fig. 2: Quantitative characterization of Distractor-phase kinematics and distraction classification model.**

**a,** The movement features that quantify responses during Distractor phase. Features of head-body angle, Δhead direction and body speed are outlined with red, yellow and blue rectangles, respectively. **b,** Comparisons of distraction metrics between distractor sides. (*N* = 10 mice). Data represent mean ± s.e.m. (black outlines). Two-tailed paired *t*-test. **c,** Proportion of distraction response levels across training sessions. Bars show the distribution of response levels pooled from all mice (*N* = 10); trial counts are indicated within the bars. **d,** Session-wise distraction metrics. DI (green) and RI (orange) are calculated from pooled trials across all mice (*N* = 10) per session. **e,** ROC curves for logistic regression models classifying distraction levels based on movement features during Distractor phase and corresponding UMAP clustering (see Fig. 1d) of the control group. The model was trained to provide an invariant classification framework for new datasets. **f,** Hierarchical clustering of Distractor-phase movement features. Clustering was performed using Spearman rank-order correlations to identify redundant variables. Dashed rectangles delineate clusters of highly correlated features defined by a linkage height threshold of 0.1. Within each cluster, a single representative feature (marked with an asterisk) was selected for modeling. **g,** Permutation importance of selected movement features for the model in **e**. Asterisks denote the representative features chosen from each correlated cluster. **h,** Coefficients of the selected movement features for the model in **e**. Coefficients for different response levels and movement feature categories are color-coded; solid and hollow bars represent positive and negative values, respectively.

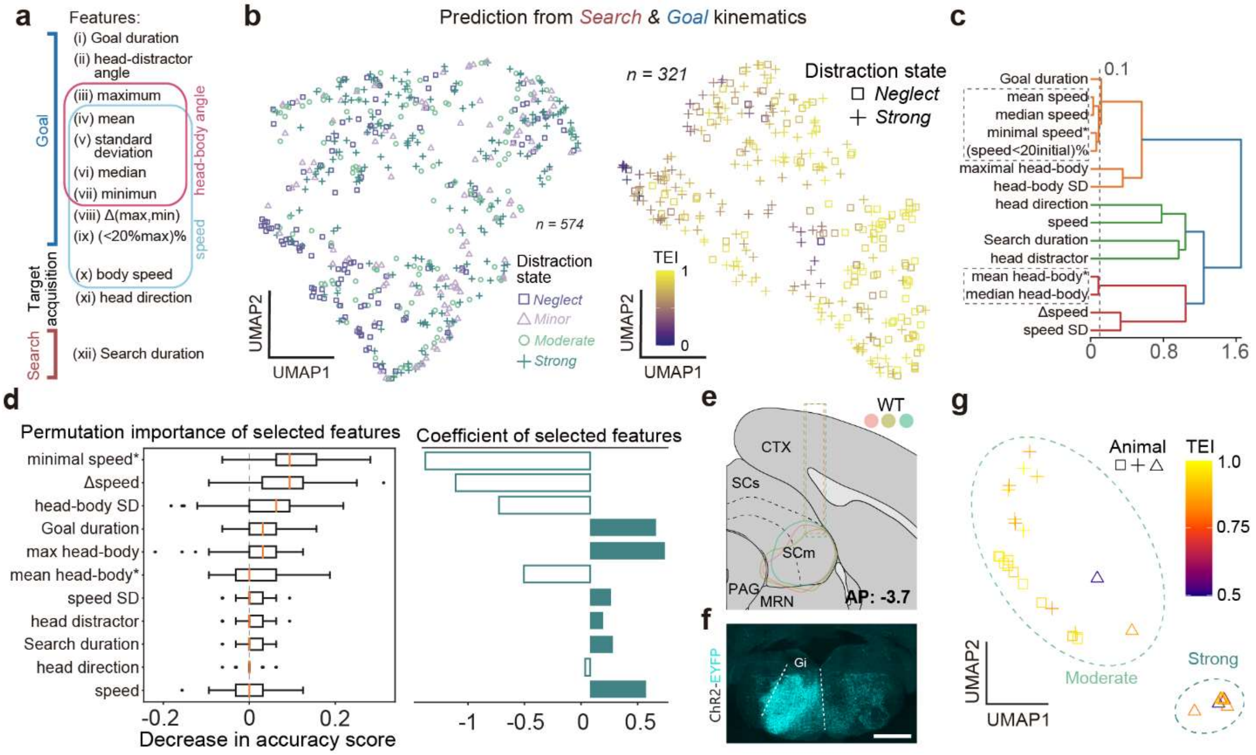


**Extended Data Fig. 3: Quantitative characterization of pre-distraction movement kinematics and the subsequent distraction prediction model**

**a,** The movement features that quantify behaviors during Search and Goal phases. Features of head-body angle and body speed are outlined with red and blue rectangles, respectively. **b,** *(Left)* UMAP embedding of pre-distraction kinematics for all trials. Each point represents a movement profile prior to distractor onset, color- and shape-coded by the subsequent response level to distractors within the same trial ($n$ = 574 responses, $N$ = 10 mice). *(Right)* Distribution of the TEI mapped onto the UMAP embedding of pre-distraction kinematics for 'Strong' and 'Neglect' trials. Color scale indicates TEI values ($n$ = 321 trials, $N$ = 10 mice). **c,** Hierarchical clustering of pre-distraction movement features based on Spearman rank-order correlation. Dashed rectangles delineate clusters of highly correlated features defined by a linkage height threshold of 0.1. Within each cluster, a single representative feature (marked with an asterisk) was selected for modeling. **d,** Permutation importance and coefficients of selected movement features for the logistic regression model in Fig. 1e predicting 'strong' versus 'neglect' trials. Asterisks denote the representative features chosen from each correlated cluster, Solid and hollow bars represent positive and negative values, respectively. **e,** Reconstruction of viral expression profiles and fiber optic implantation sites across all animals for Fig. 1g. Data from individual mice are color-coded ($N$ = 5 mice). **f,** Representative coronal section of Gi, showing the retroAAV-Cre mediated SC-Gi projections expressing ChR2-EYFP; scale bar, 1000 µm. Cyan, ChR2-EYFP. **g**, UMAP embedding of the optogenetically evoked motor responses, clusters were derived from Fig. 1i, color scale indicates TEI.

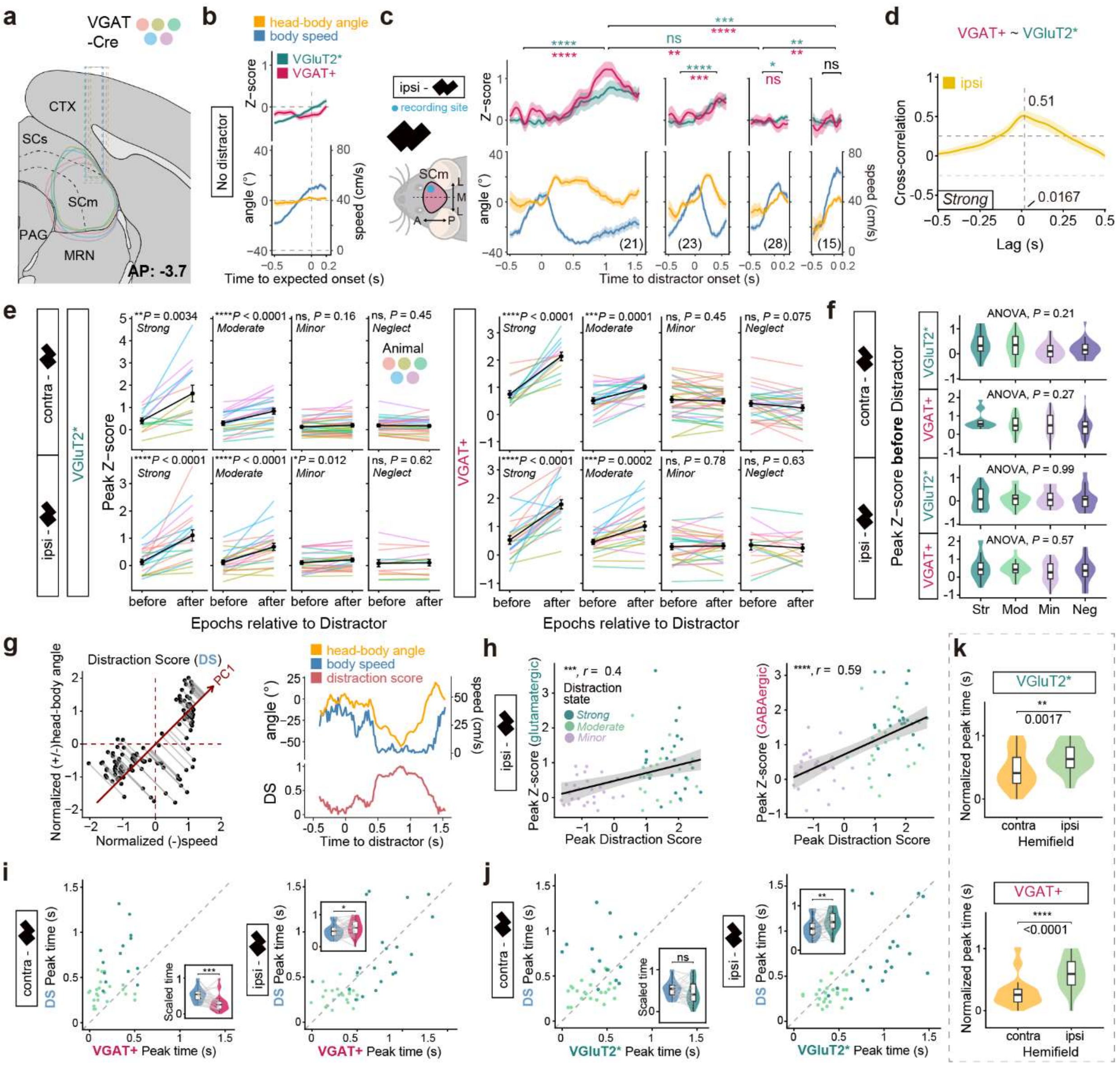


**Extended Data Fig. 4: Characterization of SC neuronal dynamics and correlation with distraction responses**

**a,** Reconstruction of viral expression profiles and fiber optic implantation sites across all animals for Fig. 2a. Data from individual mice are color-coded ($N$ = 5 mice). **b,** Average time series of z-scored calcium signals (top) and kinematic variables (bottom) for no-distractor control trials. ($n$ = 105 trials). **c**, (*Left*) Schematic of contralateral distractor placement relative to the recording site. Average time series of baseline-subtracted z-scored calcium signals (*Top*) and kinematic variables (*Bottom*) for responses to ipsilateral distractors, truncated at the median response duration. Total response counts ($n$) in parentheses. ($N$ = 5 mice). Pre- and post-distractor peak comparisons (Two-tailed paired $t$-test, FDR-corrected) and cross-level post-distractor peak comparisons (Welch's ANOVA with Games-Howell test) are indicated above traces. **d**, Cross-correlation of VGluT2* ($x$) and VGAT+ ($y$) calcium signals during 'Strong' responses for ipsilateral distractors. Dashed lines, lag at maximum correlation and 95% CIs. **e,** Comparison of mean peak calcium magnitudes of SC between the pre-distractor (-0.2 to 0 s) and post-distractor periods for each cell type and distractor location. Each line represents a single response, color-coded by individual mice. Two-tailed paired *t*-test, FDR corrected. **f,** Comparison of pre-distractor peak calcium magnitudes across all distraction levels (color-coded). Welch's ANOVA test. **g,** *(Left)* Definition of the Distraction Score (DS) on one example response. Each point represents a normalized kinematic state (speed and head-body angle, directionally adjusted) at a single time point. The red line indicates the first principal component (PC1) axis from PCA; gray lines illustrate the projection of individual points onto PC1. The DS is defined as the coordinate value along this PC1 axis. *(Right)* Time series of the example trial. **h,** Pearson's correlation between peak DS and peak z-scored calcium signals for putative VGluT2+ (VGluT2*) (*left*) and VGAT+ (*right*) neurons following ipsilateral distractors. Solid lines denote linear regression fits, shaded regions represent 95% CIs. **i, j,** Comparison of the time-to-peak for DS (blue) and VGAT+ (red, **i**) for contralateral (left; ***$P = 0.00013$) and ipsilateral (right; *$P = 0.023$) distractors or VGluT2* (green, **j**) for contralateral (left; ns, $P = 0.085$) and ipsilateral (right; **$P = 0.0013$) distractors. Two-tailed paired Wilcoxon signed-rank test. **k,** Comparison of the time-to-peak for VGluT2* (*top*) and VGAT+ (*bottom*) neurons in response to ipsilateral (green) and contralateral (orange) distractors. Two-tailed Wilcoxon rank-sum test. Data are mean ± s.e.m., vertical dashed lines indicate distractor onset ($t = 0$) unless otherwise specified. *$P < 0.05$, **$P < 0.01$, ***$P < 0.001$, ****$P < 0.0001$, ns, not significant.

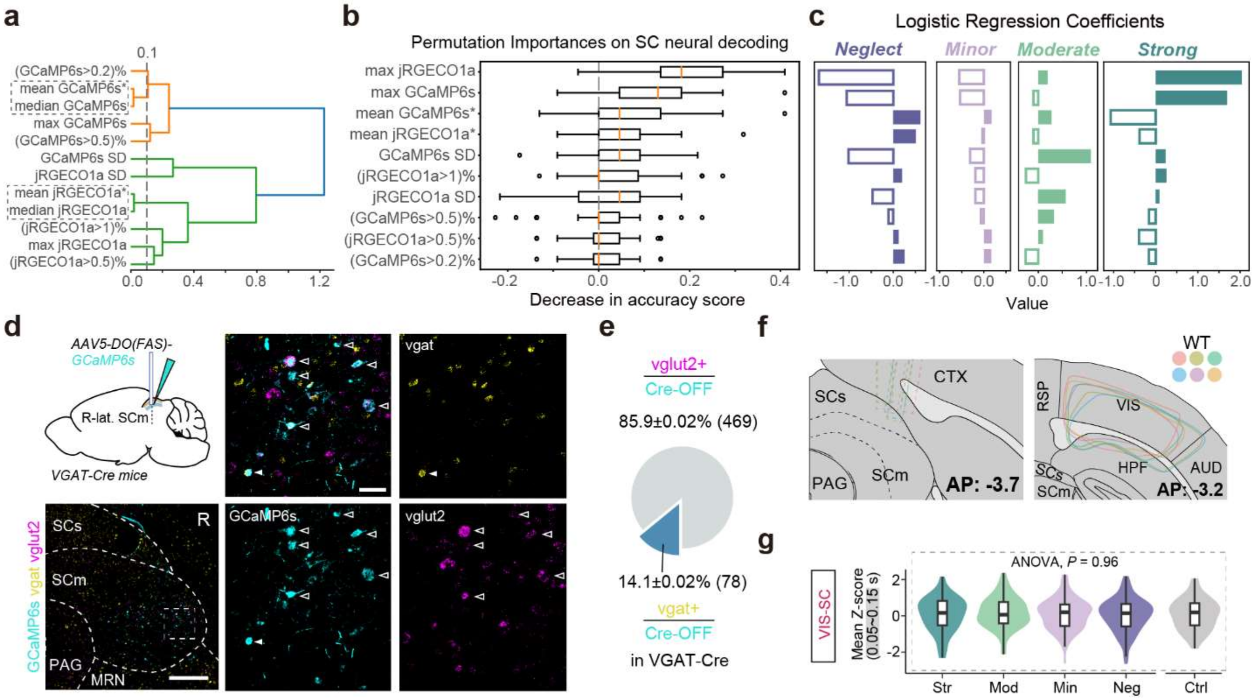


**Extended Data Fig. 5: Quantitative characterization of SC-mediated distraction prediction and technical verification of SC recording strategies.**

**a,** Hierarchical clustering of SC calcium dynamics features based on Spearman rank-order correlation. Dashed rectangles delineate clusters of highly correlated features defined by a linkage height threshold of 0.1. Within each cluster, a single representative feature (marked with an asterisk) was selected for modeling. **b, c,** Permutation importance and coefficients of selected movement features for the logistic regression model encoding all trials (in Fig. 2e). Asterisks denote the representative features chosen from each correlated cluster. **d,** (*Top Left*) Schematic of viral injection in the right lSCm for identifying neurons expressing Cre-OFF-dependent GCaMP6s in vGAT-Cre mice. (*Bottom Left*) Representative histology of a coronal brain slice of the SC showing *in situ* hybridization of *vGAT* (*Slc32a1*) and *vGlut2* (*Slc17a6*) probes along with GCaMP6s immunostaining. Scale bar, 400 µm. (*Right*) Enlarged view of the region indicated by the dashed rectangle in the bottom-*Left* panel. Solid arrowheads indicate *vGAT+*/ GCaMP6s(Cre-OFF)+ neurons; open arrowheads indicate *vGlut2+*/GCaMP6s(Cre-OFF)+ neurons. Scale bar, 50µm. Cyan, GCaMP6s; yellow, *vGAT*, magenta, *vGlut2*. **e,** Average proportion of *vGAT+* and *vGlut2+* neurons among the GCaMP6s(Cre-OFF)-labeled population (*N* = 4 mice). **f,** Reconstruction of viral expression profiles and fiber optic implantation sites across all animals for Fig. 3b. Data from individual mice are color-coded (*N* = 6 mice). **g,** Comparison of mean VIS-SC immediate responses during the shaded window across distraction levels. Welch's ANOVA test.

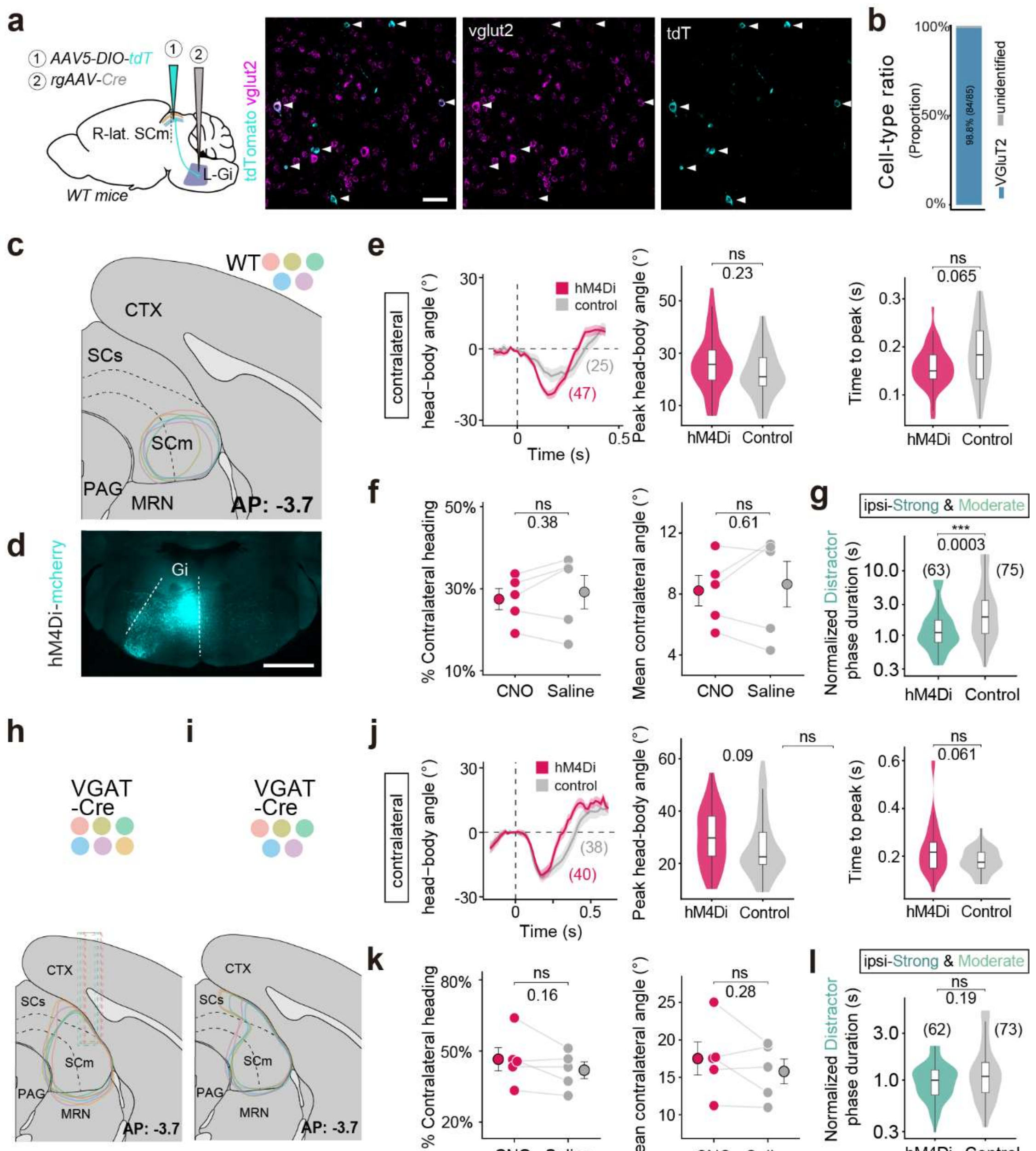


**Extended Data Fig. 6: Histological and behavioral validation for cell-type-specific SC circuit manipulations.**

**a,** (*Left*) Schematic of viral injection in the right lSCm for Gi-projecting orienting neurons expressing tdTomato in WT mice. (*Right*) Representative coronal sections of the lSCm (merged and individual channels) showing *in situ* hybridization of *vGlut2* probes along with RFP immunostaining. Solid arrowheads indicate *vGlut2+*/tdTomato+ neurons. Scale bar, 50µm. Cyan, tdTomato; magenta, *vGlut2*. **b,** Quantification of tdTomato+ cell identity. vGluT2+ (blue, *n* = 84), unidentified (grey, *n* = 1). (*n* = 2-3 sections per animal, *N* = 3 animals). **c, h, i,** Reconstruction of viral expression profiles and fiber optic implantation sites across all animals for Fig. 2f (**c**), 2i (**h**), 2n (**i**). Data from individual mice are color-coded (**c**, *N* = 5 mice, **h**, *N* = 6 mice, **i**, *N* = 5 mice). **d,** Representative coronal sections of Gi for Fig. 2f, showing the retroAAV-Cre mediated

SC-Gi projections expressing hM4Di-mCherry; scale bar, 1000 µm. Cyan, hM4Di-mCherry. **e, j,** (*Left*) Average time series of single head-turns toward contralateral distractors, representing a subset of moderate (mostly), minor, and strong responses. Data are mean ± s.e.m., vertical dashed lines indicate distractor onset (t = 0), trial counts (*n*) are in parentheses. (*Right*) comparison of peak head-body angle and time to peak between groups. Two-tailed Wilcoxon rank-sum test. **f, k,** Comparison of contralateral direction preference (defined as leftward turning >10°) and mean heading angles during task between CNO and saline sessions within the experimental group (*N* = 5 mice). Mean ± s.e.m are black-outlined with error bars. Two-tailed paired *t*-test. **g, l,** Comparison of normalized distractor phase durations of ipsilateral strong and moderate distraction responses between groups. Durations were normalized to the median values of each cluster before combining. Trial counts (*n*) are in parentheses. Two-tailed Wilcoxon rank-sum test. *P* values and significance levels are indicated.

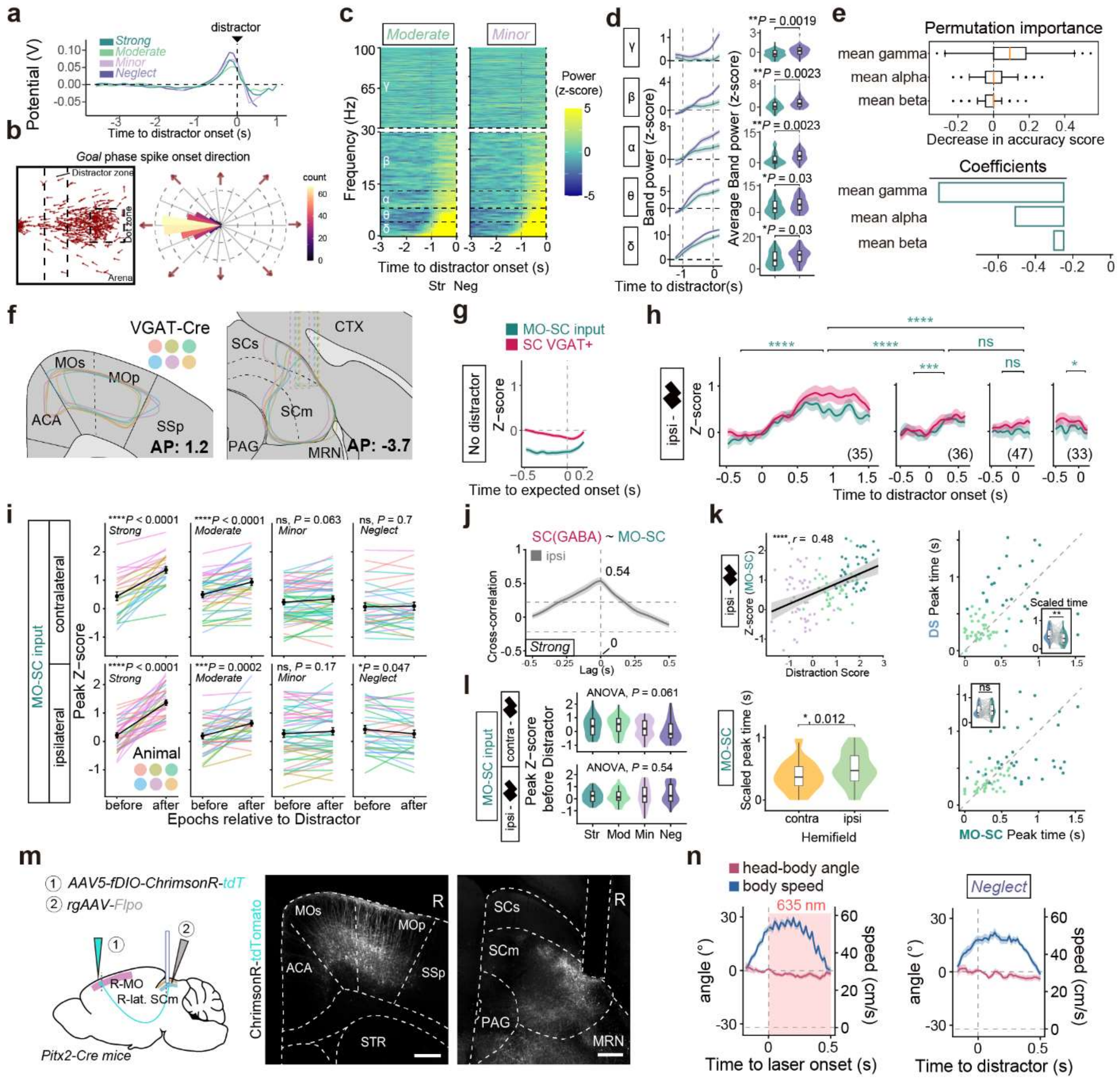


**Extended Data Fig. 7: Characterization of MO–SC projection dynamics and its quantitative role in distraction prediction.**

**a,** Average peri-distractor ECoG potentials for each distraction level (color-coded). **b,** (*Left*) Spatial distribution map of head direction vectors at spike onsets within the arena during the Goal phase. Dashed regions delineate the approximate locations of the target and distractor. Arrows indicate the instantaneous head direction. (*Right*) Polar histogram showing the angular distribution of head direction vectors. **c,** Average pre-distractor z-scored PSD or "Moderate" ($n$ = 63 trials) and "Minor" ($n$ = 58 trials) responses ($N$ = 7 mice). Dashed lines delineate pre-distractor window (-1 to 0 s) and frequency bands. **d,** (*Left*) Average z-scored power for indicated frequency bands between 'Strong' and 'Neglect'. (*Right*) Comparison of mean pre-distractor power across levels. Two-tailed $t$-test. **e,** Permutation importance (*top*) and coefficients (*bottom*) of selected band power features for the logistic regression model predicting 'Strong' and 'Neglect' trials in Fig. 4c. **f,** Reconstruction of viral expression profiles and fiber optic implantation sites across all animals for Fig. 4d. Data from individual mice are color-coded ($N$ = 6 mice). **g,** Average time series of kinematic variables (*top*) and z-scored calcium signals (*bottom*) for no-distractor control trials. Vertical dashed lines indicate simulated "distractor" onset ($t$ = 0 s) ($n$ = 250 trials). **h,** Average time series of baseline-subtracted z-scored calcium signals for responses to ipsilateral distractors, truncated at the median response duration. Total response counts ($n$) in parentheses. ($N$ = 5 mice). Pre- and post-distractor peak comparisons (Two-tailed paired $t$-test, FDR-corrected) and cross-level post-distractor peak comparisons (Welch's ANOVA with Games-Howell test) are indicated above traces. **i,** Comparison of mean peak calcium magnitudes of MO-SC inputs between the pre-distractor (-0.2 to 0 s) and post-distractor periods. Each line represents a single response, color-coded by individual mice. Two-tailed paired $t$-test, FDR corrected. **j**, Cross-correlation of SC VGAT+ ($x$) and MO-SC ($y$) calcium signals during 'Strong' responses for ipsilateral distractors. Dashed lines, lag at maximum correlation and 95% CIs. **k,** (*Top left*) Pearson's correlation between peak DS and peak z-scored calcium signals for MO-SC inputs following ipsilateral distractors. Solid lines denote linear regression fits, shaded regions represent 95% CIs. (*Bottom left*) Comparison of the time-to-peak for MO-SC projections in response to distractors. Two-tailed Wilcoxon rank-sum test. (*Right*) Comparison of the time-to-peak for DS (blue) and MO-SC projections (green) for contralateral (*top*; $^{**}P$ = 0.0027) and ipsilateral (*bottom*; ns, $P$ = 0.63) distractors. Two-tailed paired Wilcoxon signed-rank test. **l,** Comparison of pre-distractor peak calcium magnitudes across all distraction levels (color-coded). Welch's ANOVA test. **m,** (*Left*) Schematic of viral injection and fiber optic implantation in right MO and lSCm for opto-activation of MO-SC inputs in Pitx2-Cre mice. (*Right*) Representative coronal sections of MO and SC; scale bar, 400 µm. Cyan, ChrimsonR-tdTomato. **n,** Average time series of head-body angle (red) and body speed (blue) during (*Left*) laser stimulation of MO-SC projection terminals in the lSCm during the Goal phase ($n$ = 50 opto-stimulations, $N$ = 5 mice) or (*Right*) Neglect responses to distractors ($n$ = 30 responses, $N$ = 5 mice). Light blue shading indicates 635-nm laser stimulation. Data are mean ± s.e.m., vertical dashed line indicates laser or distractor onset ($t$ = 0).

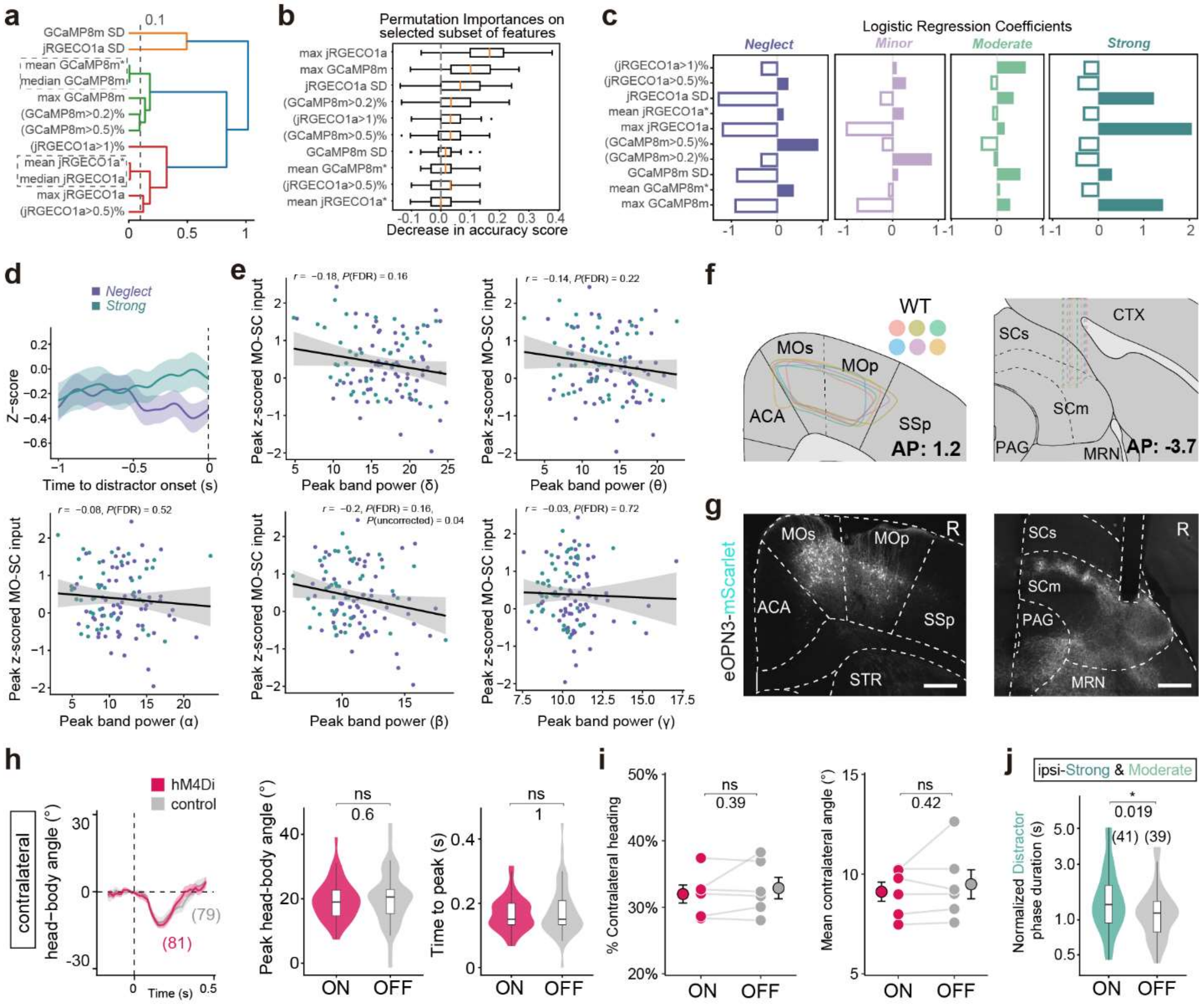


**Extended Data Fig. 8: Characterization of MO-SC encoding of visual distraction and MO–SC modulation.** **a,** Hierarchical clustering of MO-SC calcium dynamics features based on Spearman rank-order correlation. Dashed rectangles delineate clusters of highly correlated features defined by a linkage height threshold of 0.1. Within each cluster, a single representative feature (marked with an asterisk) was selected for modeling. **b, c,** Permutation importance and coefficients of selected movement features for the logistic regression model predicting all trials in Fig. 4g. Asterisks denote the representative features chosen from each correlated cluster. **d,** MO-SC dynamics before "Strong" and "Neglect" distraction responses. **e,** Correlation analysis of peak values between MO-SC dynamics and each MO band power during pre-distration period (-0.5~0 s). **f,** Reconstruction of viral expression profiles and fiber optic implantation sites across all animals for Fig. 4i. Data from individual mice are color-coded (*N* = 6 mice). **g,** Representative coronal sections of MO (left) and SC (right); scale bar, 400 µm. Cyan, eOPN3-mScarlet. **h,** (*Left*) Average time series of single head-turns toward contralateral distractors, representing a subset of moderate (mostly), minor, and strong responses. Data are mean ± s.e.m., vertical dashed lines indicate distractor onset (t = 0), trial counts (*n*) are in parentheses. (*Right*) comparison of peak head-body angle and time to peak between groups. Two-tailed Wilcoxon rank-sum test. **i,** Comparison of contralateral direction preference and mean heading angles during task between laser-ON and -OFF control sessions ($N$ = 5 mice). Mean ± s.e.m are black-outlined with error bars. Two-tailed paired $t$-test. **j,** Comparison of normalized distractor phase durations of ipsilateral strong and moderate distraction responses between groups. Durations were normalized to the median values of each cluster before combining. Trial counts ($n$) are in parentheses. Two-tailed Wilcoxon rank-sum test. $P$ values and significance levels are indicated.

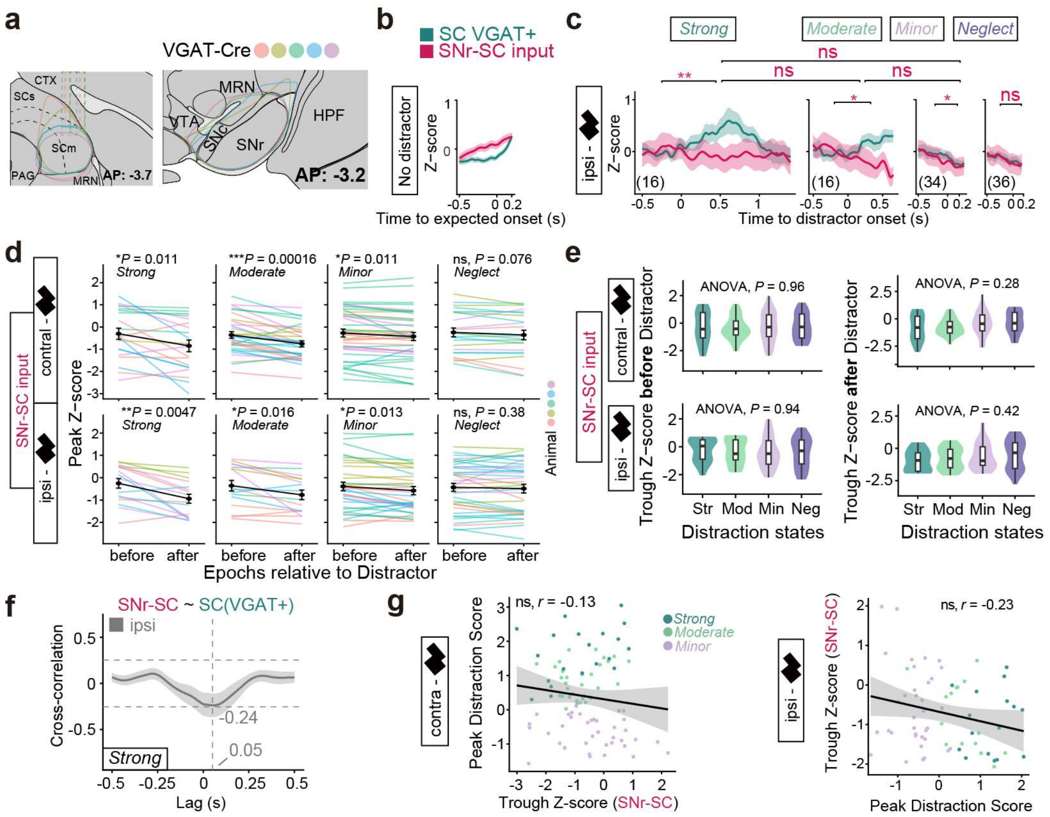


**Extended Data Fig. 9: Characterization of SNr–SC projection.**

**a,** Reconstruction of viral expression profiles and fiber optic implantation sites across all animals for Fig. 4m. Data from individual mice are color-coded (*N* = 5 mice). **b,** Average time series of z-scored calcium signals for no-distractor control trials. Vertical dashed lines indicate simulated "distractor" onset (*t* = 0 s) (*n* = 116 trials). **c,** Average time series of baseline-subtracted z-scored calcium signals for responses to ipsilateral distractors, truncated at the median response duration. Total response counts (*n*) in parentheses. (*N* = 5 mice). Pre- and post-distractor peak comparisons (Two-tailed paired *t*-test, FDR-corrected) and cross-level post-distractor peak comparisons (Welch's ANOVA with Games-Howell test) are indicated above traces. **d,** Comparison of mean peak calcium magnitudes of SNr-SC inputs between the pre-distractor (-0.2 to 0 s) and post-distractor periods. Each line represents a single response, color-coded by individual mice. Two-tailed paired *t*-test, FDR corrected. **e,** Comparison of pre- (*left*) and post- (*right*) distractor peak calcium magnitudes across all distraction levels (color-coded). Welch's ANOVA test. **f,** Cross-correlation of MO-SC (*x*) and SC VGAT+ (*y*) calcium signals during 'Strong' responses for ipsilateral distractors. Dashed lines, lag at maximum correlation and 95% CIs. **g,** Pearson's correlation between peak DS and peak z-scored calcium signals for SNr-SC inputs following contralateral (*left*) and ipsilateral (*right*) distractors. Solid lines denote linear regression fits, shaded regions represent 95% CIs.

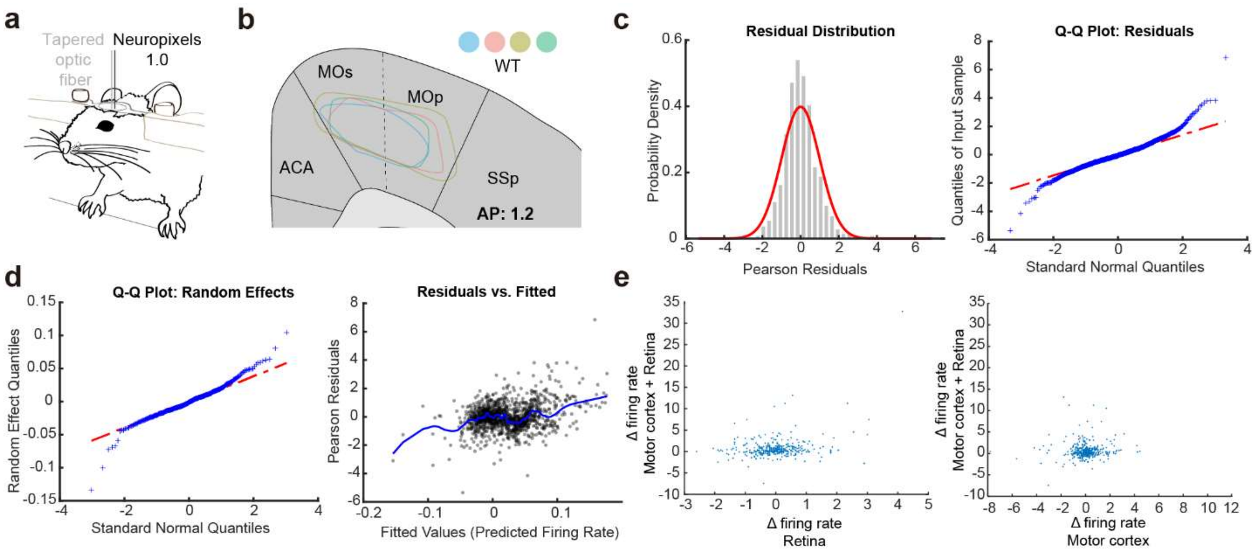


**Extended Data Fig. 10: Electrophysiological set-up, model diagnostics, and corticocollicular and retinocollicular input comparisons.**

**a**, Schematic of the head-fixed experimental configuration for electrophysiological recordings. **b,** Coronal brain atlas schematic (AP: +1.2 mm) illustrating the targeted cortical subregions, including secondary motor cortex (MOs), primary motor cortex (MOp), anterior cingulate area (ACA), and primary somatosensory cortex (SSp). Color-coded circles represent individual wild-type (WT) mice ($N = 4$). **c.** (*left)* Probability density histogram of Pearson residuals (grey) overlaid with a Gaussian fit (red curve), demonstrating a centered, symmetric residual distribution. (*Right)* Quantile-Quantile (Q-Q) plot to verify model residual normality. **d.** (*left)* Q-Q plot comparing random-effects quantiles against standard normal quantiles. (*right)* Scatter plot of Pearson residuals plotted against fitted values (predicted firing rate) from the linear mixed-effects model (LMM). **e**, Scatter plots comparing raw delta (Δ=firing rate during laser events-firing rate during baseline) rates (Hz) of individual neurons. (*left*) Simultaneous activation of cortical and retinal inputs ("Motor cortex + retina") versus retinal inputs alone ("Retina"). (*right*) Simultaneous activation ("Motor cortex + retina") versus cortical inputs alone ("Motor cortex"). Each data point represents a single neuron. Corresponding $\log_{10}$-transformed Δ firing rates are shown in Figure 6.
